# ERG orchestrates chromatin interactions to drive prostate cell fate reprogramming

**DOI:** 10.1101/2020.04.03.024349

**Authors:** Fei Li, Qiuyue Yuan, Wei Di, Xinyi Xia, Zhuang Liu, Ninghui Mao, Lin Li, Chunfeng Li, Juan He, Yunguang Li, Wangxin Guo, Xiaoyu Zhang, Yiqin Zhu, Rebiguli Aji, Shangqian Wang, Ping Chi, Brett Carver, Yong Wang, Yu Chen, Dong Gao

## Abstract

While cancer is commonly perceived as a disease of dedifferentiation, the hallmark of early stage prostate cancer is paradoxically the loss of more plastic basal cells and the abnormal proliferation of more differentiated secretory luminal cells. However, the mechanism of prostate cancer pro-luminal differentiation is largely unknown. Through integrating analysis of the transcription factors (TFs) from 806 human prostate cancers, we have identified that ERG highly correlated with prostate cancer luminal subtyping. ERG overexpression in luminal epithelial cells inhibits its normal plasticity to transdifferentiate into basal lineage and ERG supersedes PTEN-loss which favors basal differentiation. ERG knock-out disrupted prostate cell luminal differentiation, whereas AR knock-out had no such effects. Trp63 is a known master regulator of prostate basal lineage. Through analysis of 3D chromatin architecture, we found that ERG binds and inhibits the enhancer activity and chromatin looping of a Trp63 distal enhancer, thereby silencing its gene expression. Specific deletion of the distal ERG binding site resulted in the loss of ERG-mediated inhibition of basal differentiation. Thus, ERG orchestrates chromatin interactions and regulates prostate cell lineage toward pro-luminal program, as its fundamental role on lineage differentiation in prostate cancer initiation.

## Introduction

Tumor initiation, progression, and therapy resistance involve epigenetic reprogramming that leads to aberrant cell lineage specification and transition (1-5). It is critical to understand the underlying mechanisms of cancer cell lineage differentiation and transition, which will provide novel insights into anticancer research. Master transcription factors have been widely recognized with the function in cell lineage transdifferentiation and cell fate reprogramming (6-8). The identification of master transcription factors in regulation cancer cell lineage specification and transition would provide tremendous insights into the mechanism of lineage plasticity in cancer progression and therapy resistance.

Prostate cancer is the most common cancer and the second leading cause of cancer death in Western men (9). The normal prostate is a pseudostratified exocrine gland and its epithelium consists of functional luminal cells that secrete proteins of the prostatic fluid, supportive basal cells that interact with stroma and rare neuroendocrine cells. Compared to normal luminal cells, normal basal cells express higher levels of stemness genes (10) and exhibit greater stem cell properties such as increased colony and organoid formation in vitro and graft formation in vivo (11-14). While cancer is perceived as a disease of increased plasticity and stemness, primary and untreated prostate cancer is pathologically defined by luminal cell expansion and absence of basal cells with loss of P63 or CK5 by IHC. Primary and untreated prostate cancers that exhibit true basal or neuroendocrine phenotype is extremely rare (15). We and others have recently found that normal single luminal cells when grown as organoids *in vitro* or grafted *in vivo* can form normal prostate glandular structures with both secretory luminal cells and basal cells that interact with stroma (11, 12, 16). The fact that cancerous luminal cells in human prostate cancer cannot form basal cells but directly interact with stroma suggests that prostate tumorigenesis paradoxically involves a loss of normal plasticity.

Under intense selection pressure of androgen deprivation therapy that target the luminal lineage, progression to castration-resistant prostate cancer (CRPC) is associated with secondary gain of plasticity, with subsets of cancers that become neuroendocrine prostate cancer (NEPC), AR/neuroendocrine double negative prostate cancer (DNPC), AR/neuroendocrine double positive “amphicrine” prostate cancer some of which gain expression of basal markers(10, 17-19). These findings indicate that lineage differentiation and transition may play a pivotal role across multiple stages of prostate cancer progressions.

Identification on the master transcription factors has provided significant insights to understand both plasticity of prostate cancer lineages and mechanism of therapy resistances. For example, N-Myc was identified as an oncogenic driver to promote neuroendocrine prostate cancer differentiation in the context of PI3K pathway activation in both GEM mouse models (20) and transformation cellular models of human prostate epithelial cells (21). In addition, SOX2 was recognized as a key transcription factor to facilitate the lineage transitions from prostate luminal cell lineage to neuroendocrine and basal cell lineage in TP53-deficient and RB1-deficient GEM mouse models (22) as well as cellular models of human prostate cancer cell lines (23). Together, these findings proposed that SOX2 played a vital and context-dependent role for regulation on prostate cancer lineages. SOX11, as another member of SOX gene family, also promoted neuroendocrine differentiation and the treatment resistance to prostate cancers in the context of PTEN and TP53 inactivation (24). Given that the advanced prostate cancer lineage is predominantly regulated by these known transcription factors, it is reasonable to question that how primary prostate cancers gain their luminal differentiation features.

Several ETS family members have been demonstrated to be master transcription factors in the differentiation and lineage transition of several cell types (25-29). Previously, TMPRSS2-ERG fusion was reported as an early genetic alteration event occurring in 50% of prostate cancers (30, 31). Numerous previous studies revealed that ERG played an oncogenic role in promoting proliferation and invasion of prostate cancer cells (32-36). Further, we and others have shown that ERG is a master regulator that alters the chromatin enhancer landscape and AR cistrome (37-39) However, the function of ERG fusion during prostate cell lineage differentiation and transition is still largely unknown. Here, through integrating analysis of integrative classifier (40) and PAM50 classifier (41, 42), we have identified ERG with the potential role as a master transcription factor in prostate luminal lineage differentiation. In order to consolidate the further function verifications and to elucidate the detailed mechanisms, we performed multi-omics analysis (RNA-seq, ATAC-seq, ChIP-seq and 3D genome analysis BL-Hi-C) and pre-clinical model assays (*in vitro* organoids culture, *in vivo* transplantation and genetically engineered mouse models), and defined ERG as a master transcription factor to regulate prostate luminal lineage through orchestrating chromatin interactions.

## Results

### Identification for the potential master transcription factors that regulate prostate cancer cell lineage

To identify the master TFs involved in prostate cancer lineage regulation, we developed a pipeline analysis by evaluating the correlation of transcription factors with prostate cancer subtyping (Figure 1A). We firstly chose three prostate cancer cohorts with available transcriptomic profiles and each cohort contains more than 100 human prostate cancer samples (158 samples in FHCRC, 150 samples in MSKCC and 498 samples in TCGA) (31, 43, 44). In each cohort, we applied two prostate cancer subtyping methods, the PAM50 classifier categorizing prostate cancer into three lineage-related subtypes based on prostate lineage genes expression (41, 42), and the integrative classifier also revealed three distinct prostate cancer subtypes by combining several data types including transcriptomic profiles and histone modifications (40). As expected, prostate cancer samples in MSKCC cohort were categorized into three subtypes by PAM50 classifier (Supplemental Figure 1A) as well as three subtypes by integrative classifier respectively (Supplemental Figure 1B). To estimate the relationship of each TF expression levels with the above three cancer subtypes by PAM50 or integrative classifier respectively, cancer samples were further classified into three groups according to expression levels of each TF, termed as TF-high, TF-medium and TF-low. We next performed Pearson’s Chi-squared test to identify TFs that significantly correlated to prostate cancer lineages by PAM50 classifier and integrative classifier respectively. For each cohort, overlapped TFs were further defined by overlapping the identified TFs by two subtyping methods (122 in FHCRC, 208 in MSKCC and 399 in TCGA) (Supplemental Figure 1, C, D and E and Supplemental Table 1). This combinational analysis ensured that the identified overlapped-TFs would highly correlate with both prostate cancer lineages and epigenetic classifiers. Taken the reproducibility and confidence into consideration, we finally defined the 154 master TFs from those overlapped TFs that were included in at least two of the three cohorts (Figure 1B and Supplemental Table 2). Among these transcription factors, ERG showed consistent and high correlation with prostate cancer subtyping in all of the three cohorts (Figure 1C). Gene Set Enrichment Analysis (GSEA) revealed that prostate luminal cell signature (10) was significantly enriched in ERG-high prostate cancer samples, validated in two different prostate cancer cohorts (Figure 1, D and E). As expected, AR and FOXA1 were also included in these transcription factors. FOXA1 was reported with a pioneering function in prostate cancer lineage differentiation and the determination programs (45). These results revealed the high efficiency of our method to identify the master transcription factors. Overall, these results indicated that ERG, as a master transcription factor, highly correlates with prostate luminal cell lineage differentiation.

**Figure 1.**
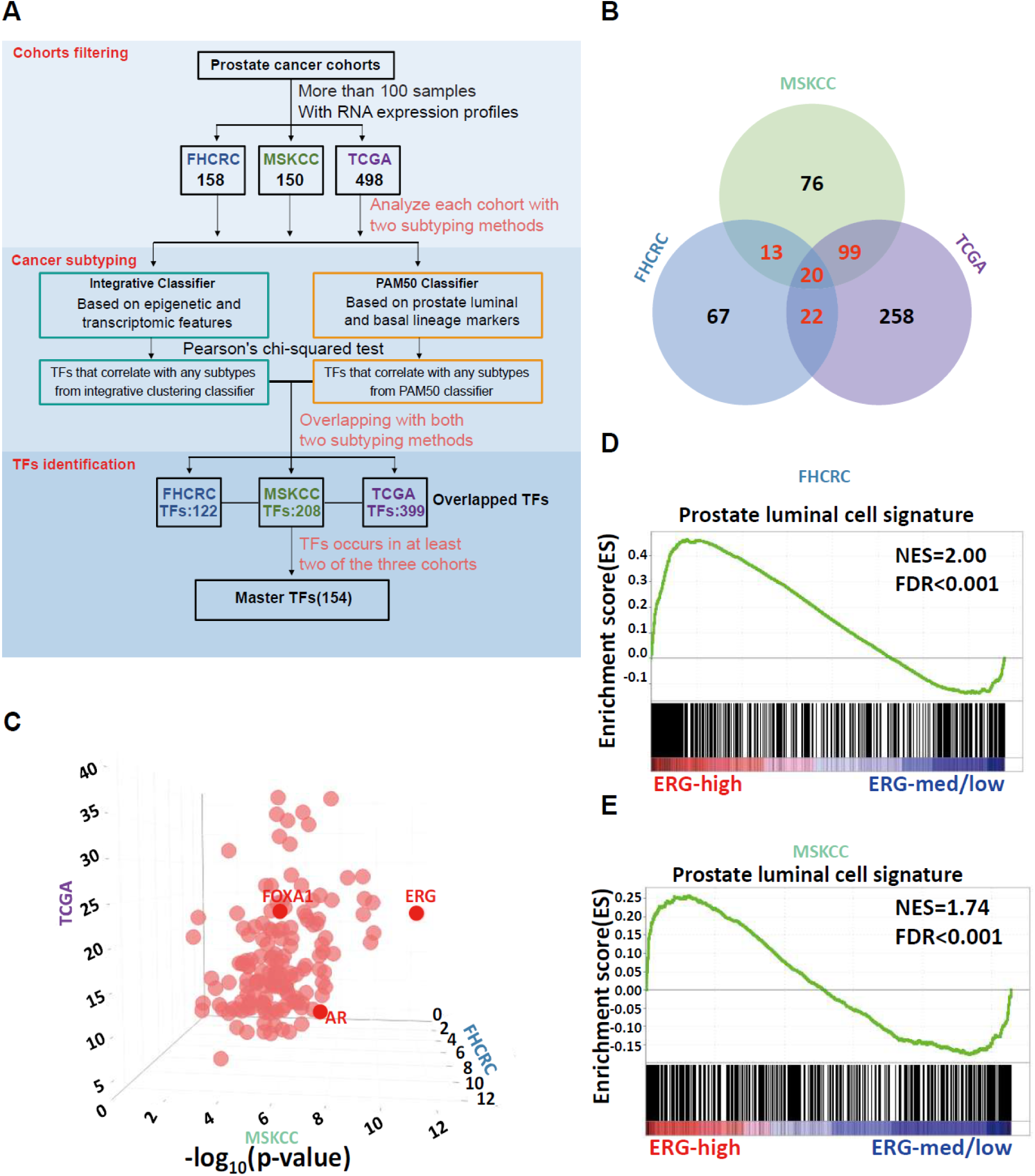
Identification for the master transcription factors that have the potential to regulate prostate cancer lineage. **(A)** Analysis pipeline to identify the master transcription factors involved in prostate cancer lineage differentiation containing three steps (1) cohorts filtering to select prostate cancer cohorts for downstream analysis, (2) cancer subtyping to categorize prostate cancer samples into several subtypes by two subtyping methods respectively and (3) TFs identification to define master TFs with high reproducibility and confidence. **(B)** Venn diagram showing the number of master TFs generated from overlapped TFs that occurs in at least two of the three cohorts (13 of MSKCC and FHCRC, 99 of MSKCC and TCGA, 22 of TCGA and FHCRC, 20 of all the three cohorts). **(C)** Bubble plot of the 154 master TFs. The value for 3 axes represents −log_10_(p-value) calculated from Pearson’s chi-squared test for MSKCC (x axis), FHCRC (y axis) and TCGA (z axis) respectively. **(D-E)** GSEA enrichment plot of ERG-high samples versus ERG-medium/low samples from FHCRC cohorts **(D)** (top) and MSKCC cohorts **(E)** (bottom) using signature genes of prostate luminal cells.

### ERG regulates normal prostate epithelial cell lineage

To investigate the cell lineage plasticity of normal prostate epithelial cells, we isolated basal cells (Epcam^+^/Cd49f^high^/YFP^-^) and luminal cells (Epcam^+^/Cd49f^low^/YFP^+^) from the anterior prostate of tamoxifen-treated *Tmprss2*^*CreERT2/+*^; *Rosa26*^*EYFP/EYFP*^ (T2Y) mice and characterized their histology features of *in vitro* organoids and *in vivo* allografts (Supplemental Figure 2, A and B) (46). Immunofluorescence analysis of luminal- and basal-cell-derived mouse prostate organoids demonstrated that both were comprised of Krt8-positive inner luminal cell layers and Krt5-positive outer basal cell layers (Supplemental Figure 2C). UGSM (urogenital sinus mesenchyme) tissue recombination assay is a useful *in vivo* method for prostate development and prostate cancer research (47). Using prostate UGSM tissue recombination assay, we further verified that both basal and luminal prostate epithelial cells from T2Y mice could reconstitute grafts with normal prostate architecture with Krt8-positive luminal cell layers and Trp63-positive basal cell layers in their renal grafts (Supplemental Figure 2D). Taken together, these results revealed that both prostate luminal and basal cells maintain bi-potential plasticity under both *in vitro* organoid culture and *in vivo* renal transplantation conditions, similar as previously reported (12, 16).

To further explore the role of ERG expression in prostate cell lineage differentiation, we isolated luminal cells from the anterior prostates of tamoxifen-treated *Tmprss2*^*CreERT2/+*^; *Rosa26*^*EYFP/ERG*^ (T2YE) mice and control T2Y mice to generate prostate organoids. Luminal-cell derived YFP-positive organoids from T2YE mice expressed ERG by IHC and were comprised of a single luminal layer of Krt8-positve cells with loss of Trp63-positive basal cells, distinct from TY mice (Figure 2A). We further analyzed the organoids derived from prostate epithelial cells of *Pb-Cre4; Rosa26*^*ERG/ERG*^ mice and *Tmprss2-ERG* knock-in mice, respectively. We confirmed that the organoids with ERG expression from these two mice also maintained luminal cell features (Supplemental Figure 2E). Next, we performed UGSM tissue recombination assays with ERG-positive and ERG-negative luminal-cell-derived organoids that were generated from T2YE and T2Y mice, respectively. The ERG-positive allografts from T2YE mice exhibited pure luminal cell features with single layer of Krt8-positive luminal cells after 2 months of transplantation (Figure 2B). On the other hand, the ERG-negative grafts from T2Y mice regenerated the normal prostate architecture composed of both Krt8-positive luminal cells and Trp63-positive basal cells (Figure 2B). Together, these results suggest that ERG overexpression could maintain the luminal cell lineage features under the conditions of both *in vitro* 3D organoid culture and *in vivo* UGSM tissue recombination.

**Figure 2.**
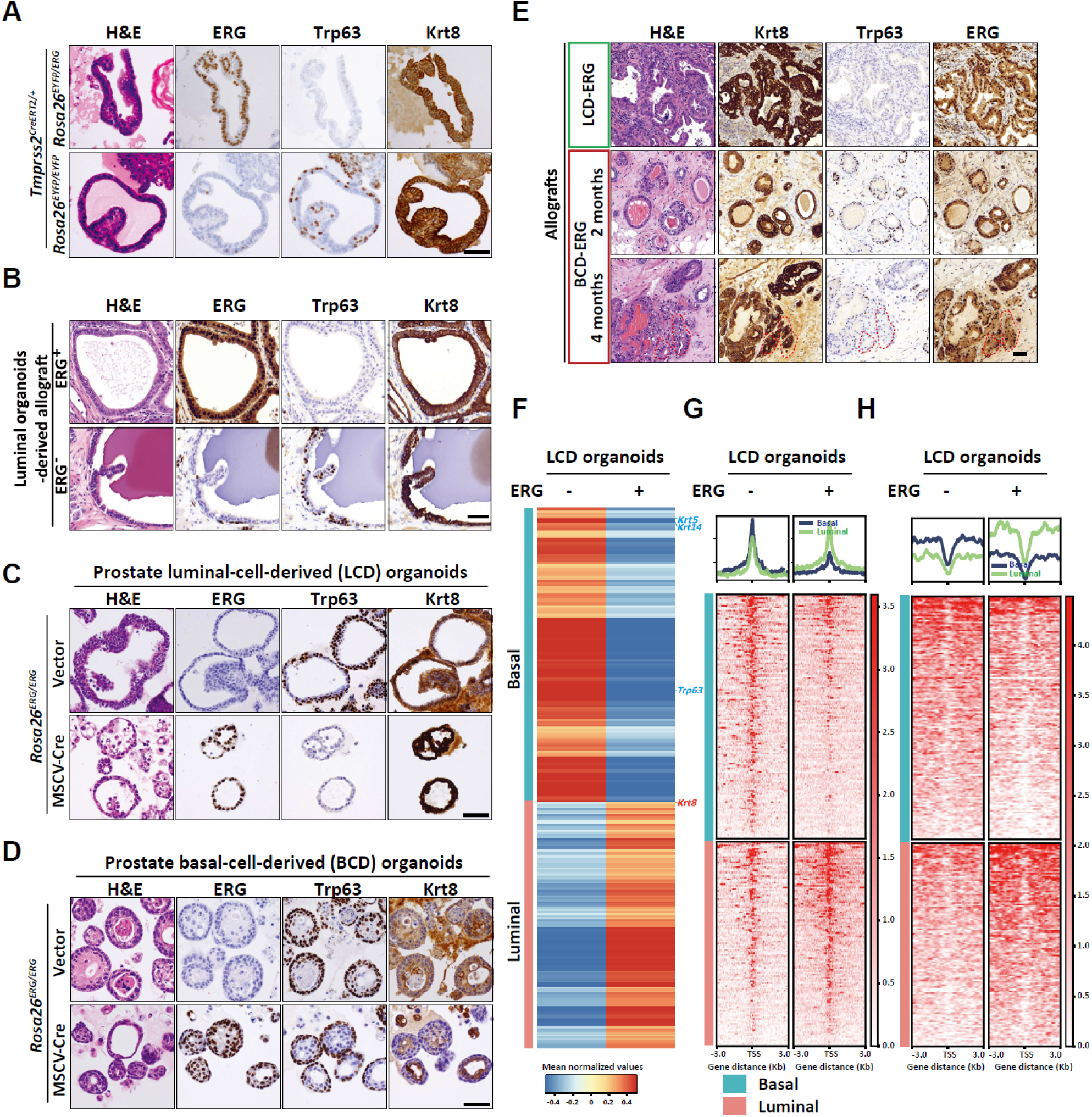
ERG promotes luminal lineage differentiation of normal prostate epithelial cells. **(A)** H&E and ERG, Trp63 and Krt8 IHC staining of luminal-cell-derived organoids generated from T2YE (top) and T2Y (bottom) mice, respectively. **(B)** H&E and ERG, Trp63 and Krt8 IHC staining of allografts derived from luminal-cell-derived organoids generated from T2YE (top) and T2Y (bottom) mice respectively. **(C-D)** H&E and ERG, Trp63 and Krt8 IHC staining of luminal-cell-derived (LCD) organoids **(C)** and basal-cell-derived (BCD) organoids **(D)** generated from *Rosa26*^*ERG/ERG*^ mice, infected with retrovirus carrying Cre recombinase (MSCV-Cre, bottom) or control backbone (MSCV-Vector, top). **(E)** H&E and Krt8, Trp63 and ERG IHC staining of allografts derived from LCD-ERG organoids (top) and BCD-ERG organoids (short term for 2 months, middle; long term for 4 months, bottom), red dashed line indicates the regions with predominant luminal phenotype. **(F)** Heatmap of RNA-seq showing the expression of down-regulated basal lineage genes and up-regulated luminal lineage genes in LCD and LCD-ERG organoids respectively. **(G-H)** Profile plot (top) and heatmap (bottom) of ATAC-seq **(G)** and H3K27ac ChIP-seq **(H)** around the transcriptional start site (TSS) of down-regulated basal lineage genes and up-regulated luminal lineage genes in LCD and LCD-ERG organoids respectively. Scale bars, 50 μm.

Given the potential role of ERG in lineage differentiation, we next sought to examine the differences of lineage responses in both luminal cells and basal cells with ERG overexpression. Briefly, we performed FACS sorting to isolate prostate luminal cells (Cd49f^low^/Cd24^high^) and basal cells (Cd49f^high^/Cd24^low^) from *Rosa26*^*ERG/ERG*^ mice (Supplemental Figure 2F). Intracellular flow cytometry for Krt5 and Krt8/Krt18 verified the cellular identities of the two populations. Next, we retrovirally transduced the Cre recombinase or a retrovirus control into these basal-cell-derived (BCD) organoids or luminal-cell-derived (LCD) organoids. Remarkably, ERG expression in luminal-cell-derived organoids (LCD-ERG) induced a single Krt8-positive luminal cell layer with loss of Trp63-positive basal cell layer, strongly indicating the predominant role of ERG in prostate cell luminal lineage differentiation (Figure 2, C and G). Basal-cell-derived organoids with ERG expression (BCD-ERG) still maintained a Trp63-positive outer basal cell layer, but with an apparent decrease in the number of Trp63-positive basal cells (Figure 2, D and G). In addition, we also performed UGSM tissue recombination assays to validate these findings *in vivo*. Allografts derived from LCD-ERG organoids exhibited pure luminal cell features with the absence of Trp63-positive basal cells after 2 months of transplantation (Figure 2E). On the other hand, allografts derived from BCD-ERG organoids were composed of both ERG^+^/Krt8^+^ luminal cells and ERG^+^/Trp63^+^ basal cells after 2 months of transplantation (Figure 2E). Intriguingly, BCD-ERG allografts also exhibited predominant luminal features with absence of Trp63-positive basal cells after 4 months of transplantation (Figure 2E). Collectively, these results demonstrated that ERG promoted prostate luminal lineage differentiation with luminal-cell-derived organoids more vulnerable to ERG-induced luminal lineage differentiation when compared to basal-cell-derived organoids.

To identify whether ERG expression-induced lineage changes would associate with chromatin status, we next performed integrative analyses of transcriptome (RNA-seq) and chromatin accessibility (ATAC-seq) of LCD organoids and LCD-ERG organoids. By assessing the lineage changes in both mRNA expression and chromatin accessibility, we identified 177 down-regulated basal signature genes, such as *Krt5, Krt14* and *Trp63*, with decreased chromatin accessibility at their promoters in LCD-ERG organoids compared to LCD organoids. On the other hand, 86 up-regulated luminal signature genes, such as *Krt8*, with increased chromatin accessibility that were identified in LCD-ERG organoids compared to LCD organoids (Figure 2, F and G). We further confirmed the increased H3K27ac levels of the up-regulated luminal signature genes and attenuated H3K27ac levels of the down-regulated basal signature genes at their promoters in LCD-ERG organoids compared to LCD organoids (Figure 2H). Collectively, these data suggested that ERG-induced changes in the expression of lineage genes were associated with chromatin status including chromatin accessibility and histone modifications, revealing the potential relationship mediated by ERG between lineage regulation and chromatin status.

### ERG regulates prostate cancer cell lineage

Given the above finding that ERG could promote luminal lineage differentiation in normal prostate epithelial cells, we next investigated that whether ERG could regulate the luminal lineage differentiation of prostate cancer cells. ERG rearrangements and loss of PTEN is regarded as one of the most concurrent genetic events in human prostate cancer (36, 48, 49). We generated *Tmprss2*^*CreERT2/+*^; *Pten*^*flox/flox*^; *Rosa26*^*ERG/ERG*^ (T2PE) mouse model to test ERG function in Pten loss condition. Due to heterogeneous recombination efficiency, ERG is only expressed in a subset of Pten deleted regions. We examined the histological features of the T2PE mice prostates at the time of 7 months after tamoxifen injection. Remarkably, ERG-positive prostate epithelial cells with Pten deletion exhibited pure Krt8-positive luminal feature and the mutual exclusions with Krt5-positive and Trp63-positive prostate basal epithelial cells (Supplemental Figure 3A). On the other hand, ERG-negative, Pten-deleted prostate epithelial cells exhibited the increased levels of basal differentiation with the expansion of Krt5-positive and Trp63-positive prostate epithelial cells (Supplemental Figure 3A). Notably, neither combination of ERG and Krt5 nor that of ERG and Trp63 showed co-localization, which were confirmed by co-staining assays of immunofluorescence (Supplemental Figure 3, B and C). Next, we isolated both *Pten*^*-/-*^ and *Pten*^*-/-*^; *R26*^*ERG*^ prostate cancer cells individually from the harvested prostates of 20-month old *Pb-Cre4; Pten*^*flox/flox*^ mice and *Pb-Cre4; Pten*^*flox/flox*^ *Rosa26*^*ERG/ERG*^ mice, respectively. We found that Pten loss prostate cancer cells differentiated towards basal lineage with the predominant expansion of Trp63-positive basal cells, while ERG overexpression maintained luminal features in the context of Pten null (Figure 3A). Compared to *Pten*^*-/-*^ organoids, the *Pten*^*-/-*^; *R26*^*ERG*^ organoids exhibited increased expression of luminal cell lineage markers, such as Krt8 and Krt18, and negative for basal cell lineage markers, such as Krt5, Krt14 and Trp63 (Supplemental Figure 3, D and E).

**Figure 3.**
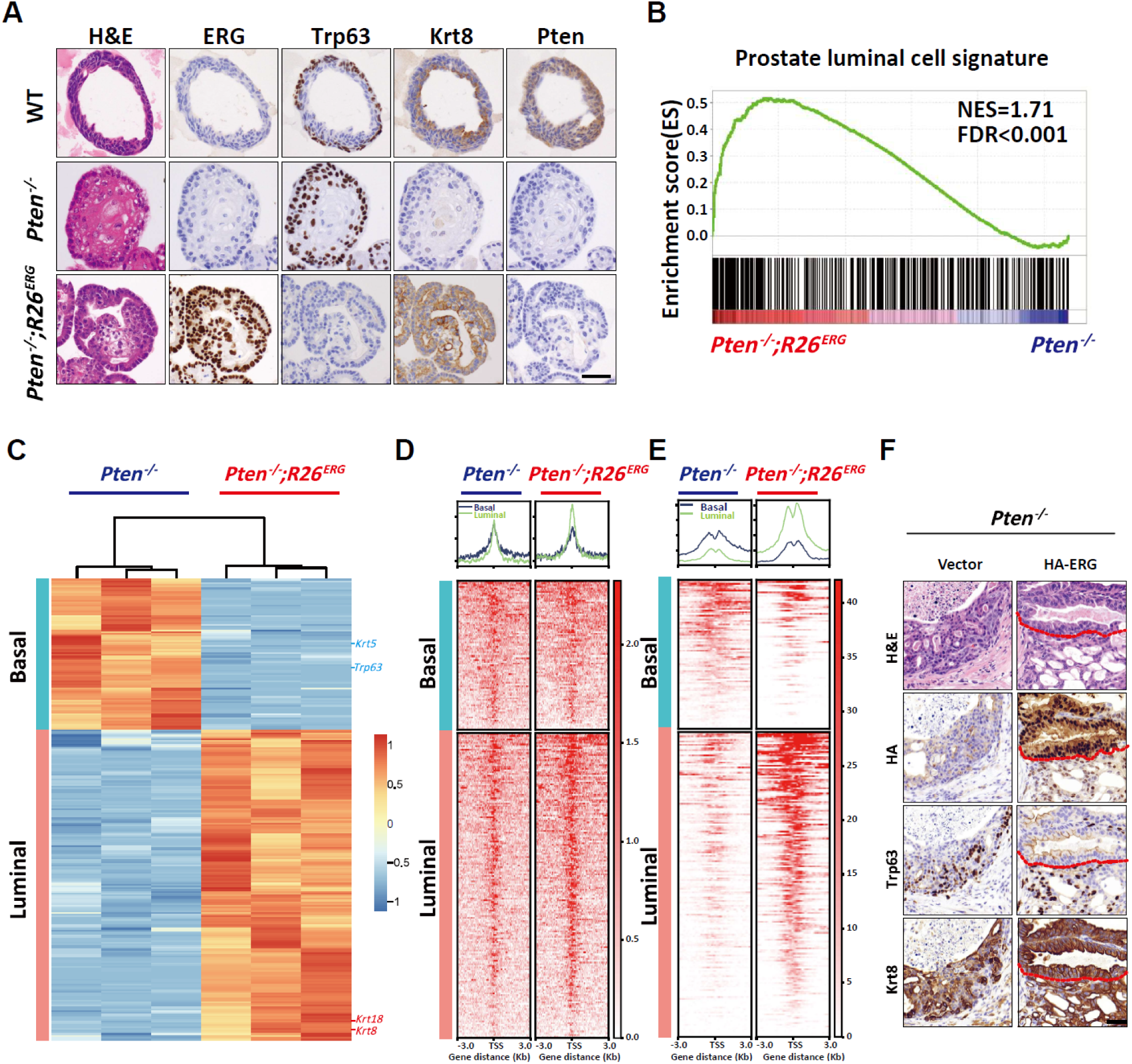
ERG promotes luminal differentiation of prostate cancer cells under the Pten loss condition. **(A)** H&E and ERG, Trp63, Krt8 and Pten IHC staining of WT (top), *Pten*^*-/-*^ (middle) and *Pten*^*-/-*^; *R26*^*ERG*^ (bottom) organoids respectively. **(B)** GSEA enrichment plot of *Pten*^*-/-*^; *R26*^*ERG*^ organoids versus *Pten*^*-/-*^ organoids using prostate luminal cell signature genes. **(C)** Heatmap showing the expression of ERG-upregulating luminal cell signature genes and ERG-downregulating basal cell signature genes in *Pten*^*-/-*^ and *Pten*^*-/-*^; *R26*^*ERG*^ organoids from RNA-seq. **(D-E)** Profile plot (top) and heatmap (bottom) of ATAC-seq **(D)** and H3K27ac ChIP-seq **(E)** around the transcriptional start site (TSS) of ERG-upregulating luminal cell signature genes and ERG-downregulating basal cell signature genes in *Pten*^*-/-*^ and *Pten*^*-/-*^; *R26*^*ERG*^ organoids respectively. **(F)** H&E and HA, Trp63 and Krt8 IHC staining of allografts derived from UGSM tissue recombination assay in SCID mice 8 weeks after transplantation of organoids overexpressing the TMPRSS2-ERG fusion protein with HA tag (right) or a control vector (left). Scale bars, 50 μm.

To characterize the impact of ERG on global gene expression, we performed RNA-seq on *Pten*^*-/-*^ and *Pten*^*-/-*^; *R26*^*ERG*^ organoids. GSEA analysis showed that the prostate luminal cell signature genes were highly enriched in *Pten*^*-/-*^; *R26*^*ERG*^ organoids rather than in *Pten*^*-/-*^ organoids (Figure 3B). Notably, the expression of luminal cell lineage markers (Krt8 and Krt18) as well as basal cell lineage markers (Krt5 and Trp63) were all included in the differentially expressed genes (DEGs) (Figure 3C), consistent with the results of qRT-PCR analyses and western blotting (Supplemental Figure 3, D and E). Here, we defined ERG-upregulating luminal signature (177 genes, Supplemental Table 3) by using the overlap between up-regulated DEGs of *Pten*^*-/-*^; *R26*^*ERG*^ organoids and prostate luminal cell signature, therefore these genes were rigorously associated with both ERG expression and prostate luminal lineage. On the other hand, ERG-downregulating basal signature (86 genes, Supplemental Table 3) was also defined by using the overlap between down-regulated DEGs of *Pten*^*-/-*^; *R26*^*ERG*^ organoids and prostate basal cell signature. Furthermore, ATAC-seq and H3K27ac ChIP-seq were performed to systematically investigate the transcriptomic and epigenetic regulations associated with ERG expression. Consistently, through ATAC-seq and H3K27ac ChIP-seq analyses, we identified the increases of both chromatin accessibility and H3K27ac levels at the promoters of luminal cell lineage markers (Krt8 and Krt18), as well as the decreases of chromatin accessibility and H3K27ac levels at the promoters of basal cell lineage markers (Krt5 and Krt14) in *Pten*^*-/-*^; *R26*^*ERG*^ organoids when compared to *Pten*^*-/-*^ organoids (Figure 3, D and E, Supplemental Figure 3F).

To further validate above results *in vivo*, we performed UGSM tissue recombination assays of *Pten*^*-/-*^ organoids overexpressing TMPRSS2-ERG fusion gene with HA tag or a control vector respectively. Consistent with our previous work, ERG overexpression efficiently promoted tumor growth (Supplemental Figure 3G) (36, 38). In addition, ERG expression promoted luminal differentiation of prostate cancer cells under the Pten loss condition (Figure 3F), further suggesting the conserved role of ERG in regulating prostate cell luminal features using multiple models. Collectively, these results suggested ERG as a master regulator to manipulate luminal lineage of prostate cancer cells, tightly associated with epigenetic regulation.

### ERG but not AR is sufficient to maintain luminal lineage in Pten loss prostate cancer

AR is a well-known transcription factor highly expressed in luminal prostate cells but is dispensable for Pten-loss mediated tumorigenesis in the mouse prostate (50, 51). To determine whether ERG or AR is required to maintain luminal differentiation in prostate cancer, we performed CRISPR/Cas9-mediated AR knock-out (AR-KO) and ERG knock-out (ERG-KO) in *Pten*^*-/-*^; *R26*^*ERG*^ organoids. AR targeted genes, such as *Fkbp5, Nkx3.1*, and *Mme*, were significantly decreased in *Pten*^*-/-*^; *R26*^*ERG*^ organoids with AR-KO (Supplemental Figure 4A). AR-KO in *Pten*^*-/-*^; *R26*^*ERG*^ organoids still maintained their pure prostate luminal histology (Krt8^+^/Trp63^-^) without obvious lineage changes, which was also evident *in vivo* UGSM tissue recombination assays (Figure 4, A and B and Supplemental Figure 4, B and C). On the contrary, ERG-KO in *Pten*^*-/-*^; *R26*^*ERG*^ organoids resulted in the loss of pure luminal differentiation and appearance of many cells that expressed basal lineage markers (Trp63, Krt5 and Krt14) in both 3D organoids and renal grafts (Figure 4, A and B and Supplemental Figure 4, B and C). In addition, the dramatic decrease on the percentage of Ki67 positive cells was attributable to ERG-KO, reinforcing the oncogenic role of ERG in the context of Pten loss (Figure 4B).

**Figure 4.**
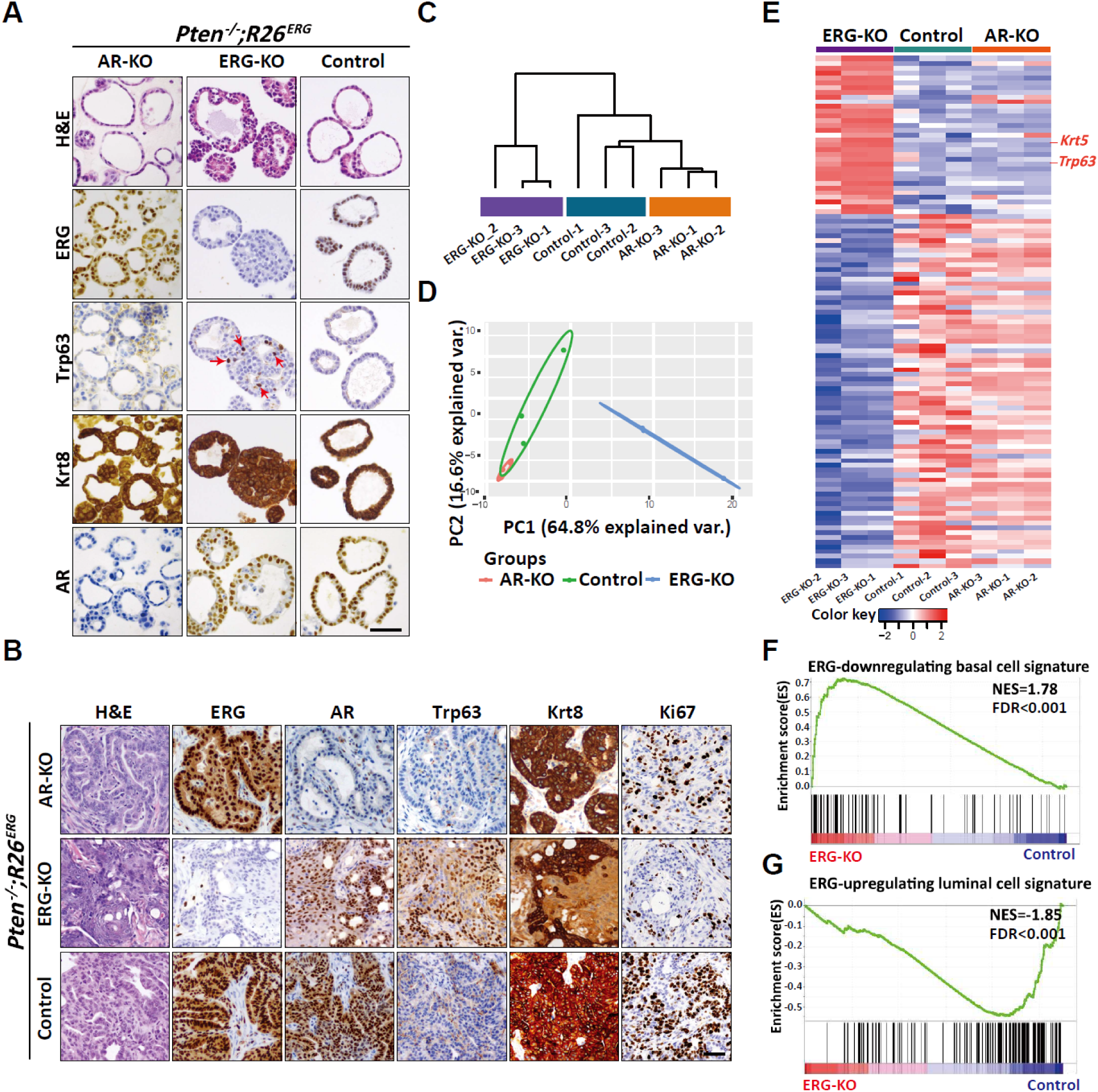
ERG not AR is required for sustaining luminal phenotype of prostate cancer cells in the context of Pten loss. **(A)** H&E and ERG, Trp63, Krt8 and AR IHC staining of *Pten*^*-/-*^; *R26*^*ERG*^ organoids infected with a lentiviral CRISPR/Cas9 carrying guide RNA targeting the AR (AR-KO, left) and ERG (ERG-KO, middle) and a control vector (Control, right), Red arrows indicate Trp63 positive cells. **(B)** H&E and ERG, AR, Trp63, Krt8 and Ki67 IHC staining of grafts derived from UGSM tissue recombination assays in SCID mice 8 weeks after transplantation of *Pten*^*-/-*^; *R26*^*ERG*^ organoids with AR-KO (top), ERG-KO (middle) and Control (bottom) respectively. **(C-D)** Clustering dendrogram **(C)** and PCA plot **(D)** for ERG-KO, AR-KO and Control organoids using prostate cell lineage signature genes. **(E)** Heatmap showing the expression of lineage-related differentially expressed genes in ERG-KO, AR-KO and Control organoids. **(F-G)** GSEA enrichment plot of ERG-KO versus Control using ERG-downregulating basal cell signature genes **(F)** and ERG-upregulating luminal cell signature genes **(G)** respectively. Scale bars, 50 μm.

Based on the expressions of prostate lineage genes, hierarchical clustering and PCA analyses were performed to evaluate the similarities among AR-KO, ERG-KO and control *Pten*^*-/-*^; *R26*^*ERG*^ organoids. AR-KO showed only relatively small changes with control *Pten*^*-/-*^; *R26*^*ERG*^ organoids, while ERG-KO organoids were clearly separated from AR-KO and control *Pten*^*-/-*^; *R26*^*ERG*^ organoids (Figure 4, C and D). Moreover, remarkable increased basal cell lineage markers, such as *Krt5* and *Trp63*, were also verified in ERG-KO organoids through RNA-seq DEGs analyses of AR-KO, ERG-KO and control *Pten*^*-/-*^; *R26*^*ERG*^ organoids (Figure 4E). Furthermore, GSEA was performed to evaluate the changes of prostate cell lineage, revealing that ERG-downregulating basal signature genes were significantly enriched in ERG-KO organoids, while ERG-upregulating luminal signature genes were enriched in control *Pten*^*-/-*^; *R26*^*ERG*^ organoids (Figure 4, F and G). Consistently, no significant differences were identified in the expression of prostate lineage genes between AR-KO and control (Supplemental Figure 4D). These findings further confirmed the importance of ERG in the lineage regulation of prostate cancer cells. Moreover, in the context of both Pten loss and ERG expression, AR deletion had no significant effects on prostate cell lineage differentiation, suggesting that luminal lineage regulation in primary prostate cancer cells does not rely on AR.

### ERG induces the global changes in chromatin interactions

Chromatin dynamics are highly correlated with cell fate reprogramming (52-54). To examine whether ERG expression induces changes in chromatin interactions, we performed Bridge Linker-Hi-C (BL-Hi-C)(55) in LCD, LCD-ERG, *Pten*^*-/-*^ and *Pten*^*-/-*^; *R26*^*ERG*^ organoids. On average, each library contained over 470 million unique pairwise contacts, which had high quality with over 80% percentage of *cis-*pairs in total valid pairs (Supplemental Figure 5, A and B). After the systematic loop calling, we found that ERG expression resulted in the increased number of interaction loops in the contexts of both Pten intact and Pten loss (Figure 5A and Supplemental Figure 5C). Circos plot to globally visualize the differential interactions (DIs) across the 21 chromosomes (chromosome 1-19, X and Y) demonstrated that ERG expression enhanced chromatin interactions (Figure 5B and Supplemental Figure 5D). To investigate the associations between chromatin interactions and gene expression, we next correlated DEGs with DIs of *Pten*^*-/-*^; *R26*^*ERG*^ organoids compared to *Pten*^*-/-*^ organoids (Figure 5C). Remarkably, the percentages of down-regulated DEGs with DIs reached to 81% (711/873, *p*=5.89e-83), including *Trp63*, and *Krt5*. Moreover, 82% (1910/2342, *p*=2.77e-176) of up-regulated DEGs were found with DIs, including *Krt8* and *Krt18*. When similar analyses were performed on both LCD-ERG and LCD organoids in the Pten-intact setting, we found that 79% (1270/1612, *p*=1.82e-118) of down-regulated DEGs as well as 80% (802/1001, *p*=4.61e-89) of up-regulated DEGs were both mapped with DIs respectively (Supplemental Figure 5E). To further explore the enrichment pattern of chromatin interactions in prostate lineage related loci, the *Krt8* and *Krt18* genomic regions in chromosome 15 were chosen with contact maps shown at 20-and 1-kb resolution. Upon close inspections on these regions, we observed that enhanced chromatin interactions were detected in ERG-expressing organoids, including both *Pten*^*-/-*^; *R26*^*ERG*^ organoids (Figure 5D) and LCD-ERG organoids (Supplemental Figure 5F). These observations indicate that the gene expression changes induced by ERG were highly associated with the alterations of chromatin interactions. To directly characterize the role of ERG in chromatin interactions, we binned the genome into 1-Mb intervals and analyzed the total DIs in these genomic bins respectively. Importantly, we observed the preferential ERG binding occupancy of genomic bins with more DIs (Figure 5E and Supplemental Figure 5G), such positive correlation was also confirmed by Pearson correlation analysis (Figure 5F and Supplemental Figure 5H). Taken together, these results suggest that ERG binding occupancy significantly correlated with differential chromatin interactions, which also highly associated with differentially expressed genes, indicating the potential role of ERG in transcriptional programs through re-organizing chromatin interactions to facilitate cell lineage regulation.

**Figure 5.**
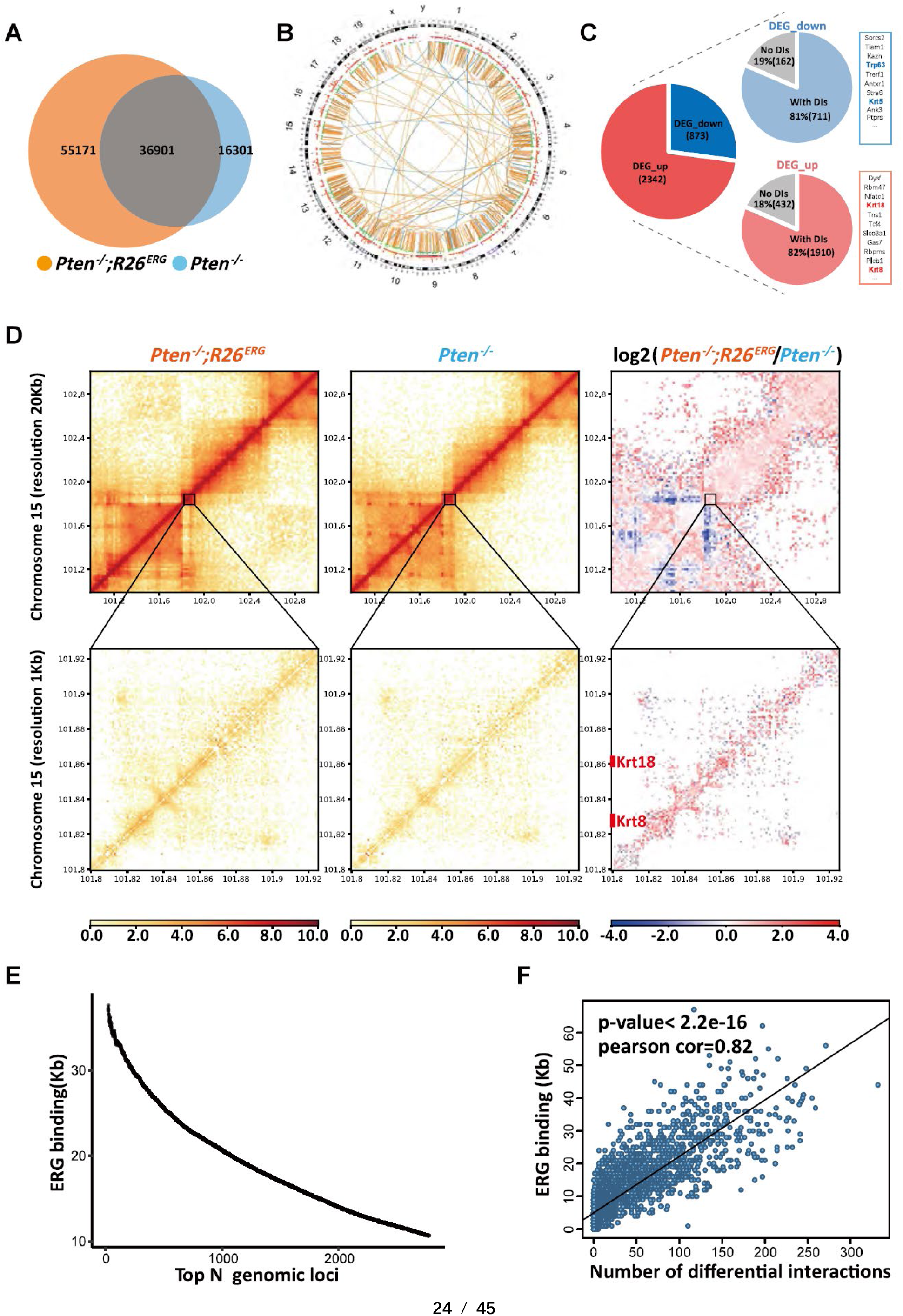
ERG globally alters chromatin interactions that are associated with gene expression changes. **(A)** Venn plot showing differential chromatin interactions between *Pten*^*-/-*^ and *Pten*^*-/-*^; *R26*^*ERG*^ organoids, orange circle and light blue circle represent chromatin interactions of *Pten*^*-/-*^; *R26*^*ERG*^ and *Pten*^*-/-*^ respectively. **(B)** Circos plot depicting chromosomes 1-19, X and Y on the basis of BL-Hi-C data and RNA-seq data, indicating DIs including *Pten*^*-/-*^; *R26*^*ERG*^ -specific DIs(orange) and *Pten*^*-/-*^-specific DIs(light blue), DEGs including up-regulated DEGs of *Pten*^*-/-*^; *R26*^*ERG*^ (red) and down-regulated DEGs of *Pten*^*-/-*^; *R26*^*ERG*^ (green), respectively. **(C)** Pie plots showing the percentage of down-regulated DEGs with DIs (top) and up-regulated DEGs with DIs (bottom). **(D)** The normalized interaction heatmaps of *Pten*^*-/-*^; *R26*^*ERG*^ (left), *Pten*^*-/-*^ (middle), and the difference (right) at 20 kb resolution (top) and 1 kb resolution (bottom) of chromosome 15, including Krt8 and Krt18 gene loci. **(E)** Plot showing the density of ERG binding (Kb) at each of the ranked (N) differential interacting chromatin loci of 1-Mb intervals. **(F)** Correlation plot showing the significant positive relationship between ERG binding density and the number of DIs in 1-Mb intervals.

### Deletion of a specific ERG binding site disrupts the function of ERG in prostate lineage regulation

Given the associations between transcriptional regulations induced by ERG and chromatin interactions, we next asked whether such associations were functionally related to prostate lineage regulation. Through integrating motif enrichment analysis with transcriptional expression changes generated from ATAC-seq and RNA-seq respectively, we found that Trp63 exhibited high potential as a master transcription factor in both LCD organoids and *Pten*^*-/-*^ organoids, both of which contained cells with basal cell differentiation (Supplemental Figure 6A). Concordantly, ERG played a pivotal role in prostate lineage regulation that was verified in both LCD-ERG organoids and *Pten*^*-/-*^; *R26*^*ERG*^ organoids (Supplemental Figure 6A). Indeed, Trp63 is a known master regulator of the prostate basal cell lineage and *Trp63* knockout mice failed to develop basal cells (56, 57).

To determine whether Trp63 expression could be regulated by ERG through altering chromatin interactions, we first examined the chromatin interactions of the *Trp63* (ΔNp63) loci in *Pten*^*-/-*^ and *Pten*^*-/-*^; *R26*^*ERG*^ organoids by BL-Hi-C. The attenuated chromatin interactions of the *Trp63* loci were identified, whereas the chromatin interactions of its neighboring gene loci, *Leprel1*, were remarkably increased in *Pten*^*-/-*^; *R26*^*ERG*^ organoids compared to *Pten*^*-/-*^ organoids (Figure 6A). In addition, almost all the chromatin interactions of the *Trp63* loci were distributed between the loci and the region at 400 kb upstream of the *Trp63* promoter in both *Pten*^*-/-*^ and *Pten*^*-/-*^; *R26*^*ERG*^ organoids. Intriguingly, this region was accompanied by a strong ERG binding site in *Pten*^*-/-*^; *R26*^*ERG*^ organoids (Figure 6A). This result indicated a potential role of this ERG binding site in mediating the associations between *Trp63* expression and chromatin interactions. We next specifically investigated the chromatin interactions and histone modifications between this distal ERG binding site and *Trp63* loci. Upon close inspections on this region, we observed an enhancer strongly enriched for the H3K27ac histone mark in ERG-negative LCD organoids and *Pten*^*-/-*^ organoids, suggesting this is a bona fide enhancer for *Trp63* in prostate cells (Figure 6B and Supplemental Figure 6B). Upon ERG expression in LCD-ERG organoids and *Pten*^*-/-*^; *R26*^*ERG*^ organoids we did not observe ERG binding to *Trp63* gene body, but to its distal enhancer (Figure 6B and Supplemental Figure 6B). Remarkably, there were significantly decreased H3K27ac signals at the distal enhancer as well as chromatin interaction loops with the Trp63 promoter upon ERG binding site in both *Pten*^*-/-*^; *R26*^*ERG*^ (Figure 6B and 6C) and LCD-ERG organoids (Supplemental Figure 6, B and C). These results indicate a functional link between ERG-directed re-wiring of chromatin interactions and epigenetic modifications to regulate gene expression.

**Figure 6.**
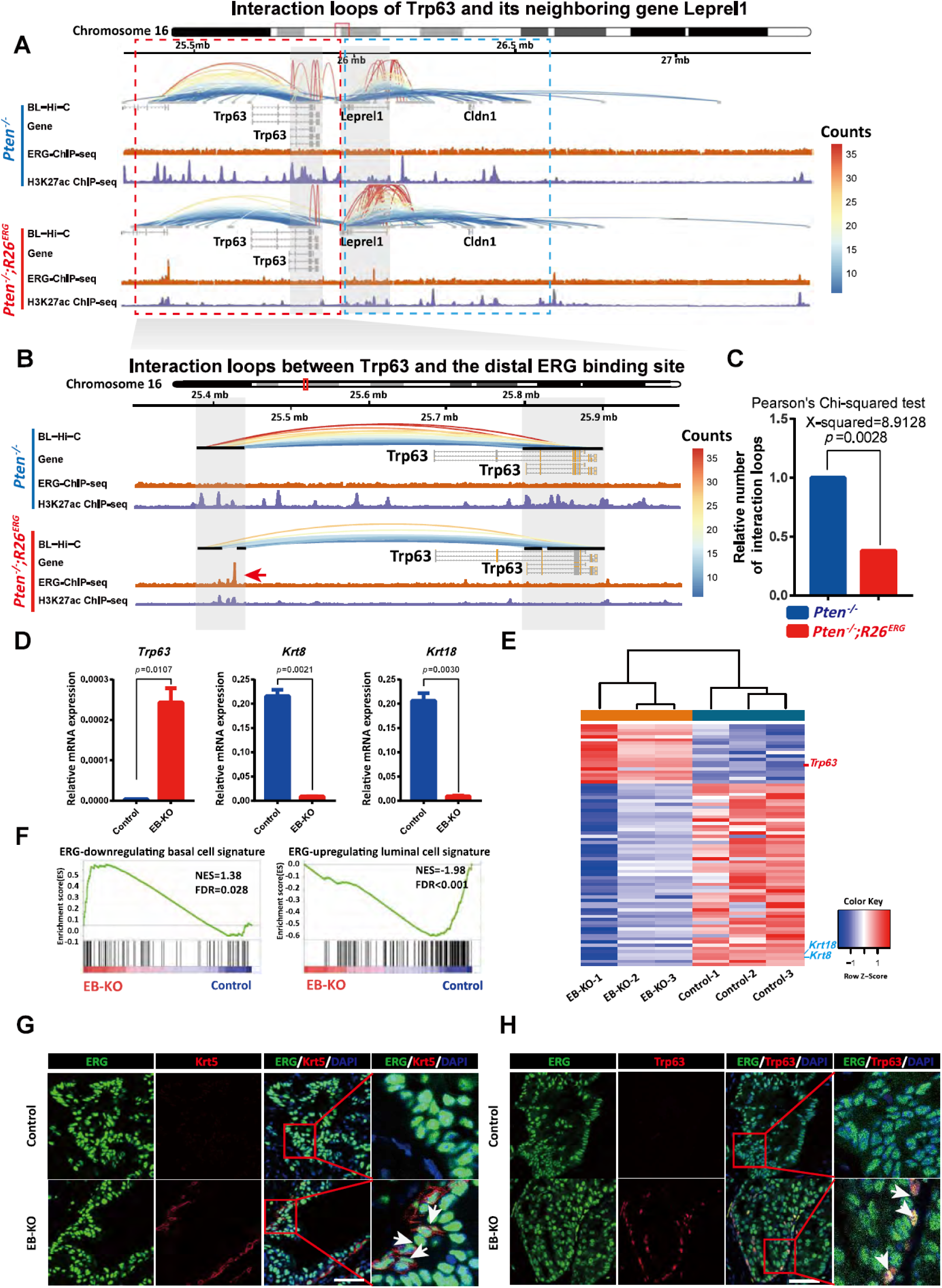
Deletion of a specific ERG binding site impaired the function of ERG in prostate lineage regulation. **(A)** 3D signal of BL-Hi-C showing chromatin interactions of *Trp63* loci and its neighboring gene *Leprel1* loci in *Pten*^*-/-*^ (top) and *Pten*^*-/-*^; *R26*^*ERG*^ (bottom) organoids respectively, red box indicates the highly interacting region of *Trp63* loci, blue box indicates the highly interacting region of *Leprel1* loci. **(B)** 3D signal of BL-Hi-C showing chromatin interactions between the distal ERG binding site and Trp63 gene body region in *Pten*^*-/-*^ (top) and *Pten*^*-/-*^; *R26*^*ERG*^ (bottom) organoids respectively. Red arrow indicates the distal ERG binding site. **(C)** Pearson’s Chi-squared test to evaluate the differences of interaction loops density between *Pten*^*-/-*^ and *Pten*^*-/-*^; *R26*^*ERG*^ organoids. **(D)** QRT-PCR analysis of *Trp63, Krt8* and *Krt18* mRNA expression in EB-KO and Control of *Pten*^*-/-*^; *R26*^*ERG*^ organoids (two-tailed t-test, mean ± sem). **(E)** Heatmap of RNA-seq for EB-KO and Control of *Pten*^*-/-*^; *R26*^*ERG*^ organoids using differentially expressed prostate cell lineage signature genes. **(F)** GSEA enrichment plot of EB-KO organoids versus Control organoids using ERG-downregulating basal cell signature genes (left) and ERG-upregulating luminal cell signature genes (right) respectively. **(G)** ERG, Krt5 and DAPI IF staining for allografts of UGSM tissue recombination assays derived from EB-KO and Control organoids, arrows indicate ERG^+^/Trp63^+^ cells. **(H)** ERG, Trp63 and DAPI IF staining of allografts from UGSM tissue recombination assays derived from EB-KO and Control organoids, arrows indicate ERG^+^/Trp63^+^ cells. Scale bars, 50 μm.

**Figure 7.**
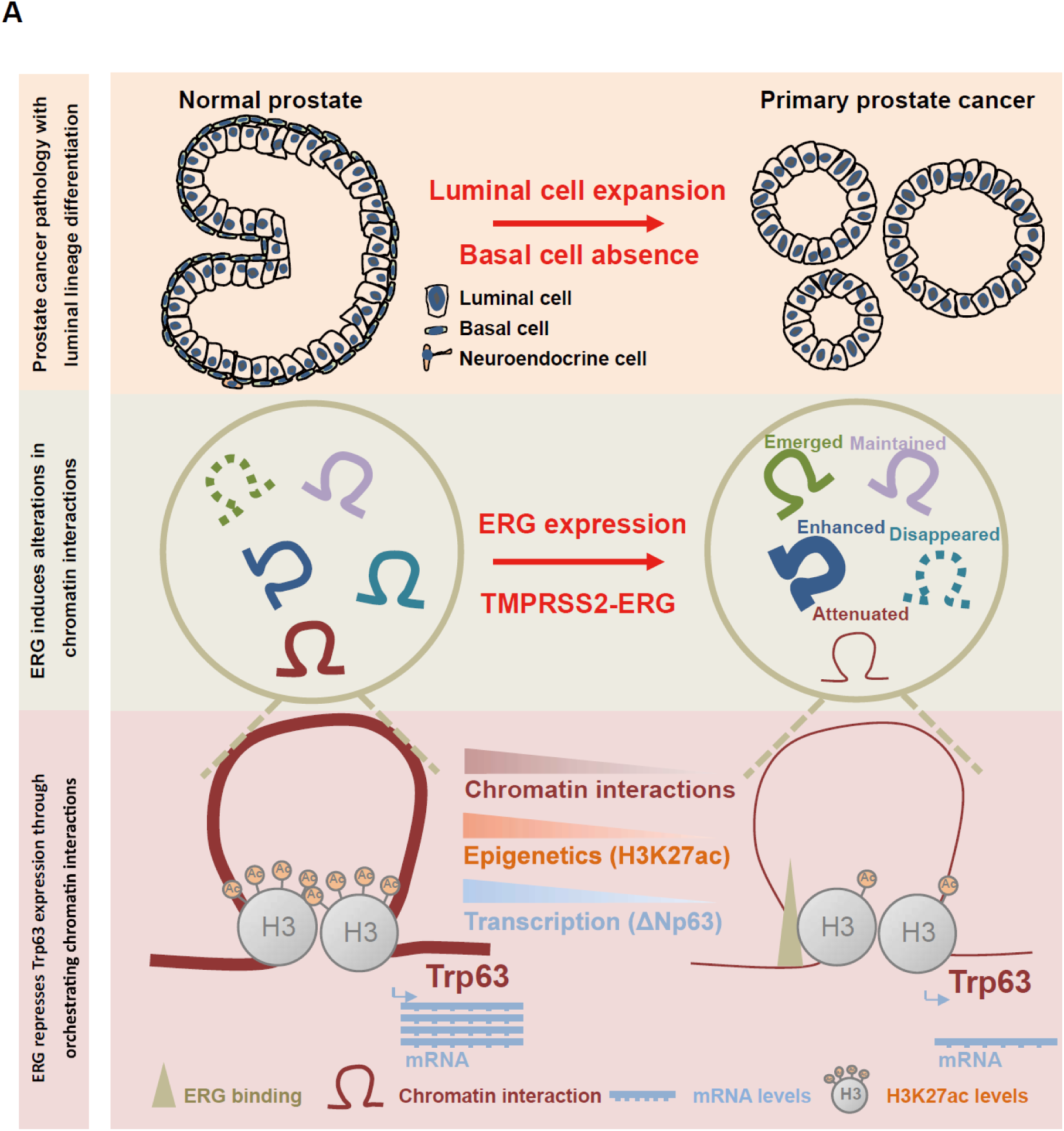
Schematic diagram for ERG drives prostate cell fate reprogramming through orchestrating chromatin interactions. **(A)** Most of prostate cancers are characterized by luminal cells expansion and basal cells absence, compared to normal prostate architecture that are composed of both luminal cells and basal cells (top). ERG overexpression driven by TMPRSS2-ERG fusion is one of the most common genetic alteration events in prostate cancer, which can alter chromatin interactions (middle). Since chromatin architecture is closely associated with epigenetic modifications and mRNA transcription, ERG-induced alterations in chromatin interactions may cause dysregulation of genes including Trp63. ERG overexpression reduces chromatin interactions and H3K27ac levels across the region from a distal ERG binding site to Trp63 gene body, which further causes decreased mRNA levels of Trp63 to facilitate the function of ERG in promoting luminal lineage differentiation (bottom).

To further determine whether ERG could directly repress Trp63 expression through the distal binding-induced attenuations on chromatin interactions, we used CRISPR/Cas9 system to specifically delete this ERG binding site in *Pten*^*-/-*^; *R26*^*ERG*^ and LCD-ERG organoids. Results of Sanger sequencing confirmed the *Pten*^*-/-*^; *R26*^*ERG*^ organoid clones with the successful ERG binding site heterozygous knock-out (Supplemental Figure 6D). Notably, both qRT-PCR and western blotting assays revealed that deletion of the ERG binding site (EB-KO) resulted in the increased expression of Trp63 and the decreased expression of both Krt8 and Krt18 in *Pten*^*-/-*^; *R26*^*ERG*^ organoids (Figure 6D and Supplemental Figure 6E) as well as LCD-ERG organoids (Supplemental Figure 6, H and I). To further characterize the global changes of prostate lineage induced by EB-KO, we compared three independent EB-KO organoid clones with control *Pten*^*-/-*^; *R26*^*ERG*^ organoid clones using ERG-regulating prostate lineage genes. GSEA demonstrated that EB-KO was significantly associated with the reduced expression of ERG-upregulating luminal signature genes and the increased expression of ERG-downregulating basal signature genes (Figure 6, E and F). Furthermore, PCA analysis revealed the distinct relationships among EB-KO, ERG-KO and control in each of their *Pten*^*-/-*^; *R26*^*ERG*^ organoids. Intriguingly, ERG control showed a closer relationship with EB-KO than that with ERG-KO, suggesting that EB-KO could partially phenocopy the biological effects of ERG-KO (Supplemental Figure 6F). Given the effects of EB-KO on lineage changes *in vitro*, we next sought to investigate the effects of EB-KO *in vivo* using UGSM tissue recombination assays. Remarkably, outer layer with Trp63^+^/ERG^+^ and Krt5^+^/ERG^+^ basal cells could be widely identified in EB-KO-derived allografts, indicating the EB-KO-induced differentiation of prostate basal lineage *in vivo* (Figure 6,G and H and Supplemental Figure 6G).

To validate the existence of the distal ERG binding site in human prostate cells, we analyzed a dataset that was previously generated from RWPE-1 cells with ERG overexpression (58). Remarkably, we found the actual existence of distal ERG binding site in ERG-expressing RWPE-1 cells (Supplemental Figure 7A). Moreover, the homologies for these binding sites in prostate cells between human and mouse were also confirmed by the additional analyses using NCBI BLAST tools (Supplemental Figure 7B). We next sought to characterize the lineage changes induced by ERG expression in human prostate cells. Consistent with those results found in mouse prostate cells, ERG expression resulted in the enhanced luminal phenotype with the increased expression of *KRT8* and *KRT18*, and attenuated the basal phenotype indicated by the reduced expression of *TP63, KRT5* and *KRT14* (Supplemental Figure 7,C, D and E).

In summary, our above results demonstrated that the function of distal ERG binding site in ERG-mediated maintenances and regulation on prostate luminal cell features, reflecting that ERG orchestrates the plasticity of prostate luminal lineage through chromatin interactions. In addition, the existence of the distal ERG binding site in human prostate cells reveals a conserved role of ERG in prostate luminal lineage regulation.

## Discussion

Definitive evidence collected during past years supports the close associations between activity of transcription factors (TFs) and cell lineage determination in various biological processes, including development, immune response and cancer progression (59-62). Particularly, primary prostate cancer is characterized with both luminal cells expansion and loss of basal cells. Therapeutic treatments on prostate cancers can select for lineage alterations with the transitions from luminal cell lineage toward neuroendocrine and basal differentiation. Numerous studies have focused on lineage transitions in CRPC. However, the lineage determining mechanism of primary prostate luminal cancers are still largely unknown. Here, we have successfully identified ERG as a master regulator in regulating prostate cancer cell luminal lineage through chromatin interaction changes.

TMPRSS2-ERG fusion is a common genetic alteration event (∼50%), which drives ERG expression occurring in the early-stage of prostate cancer (30). We identified ERG as a master regulator in prostate cancer lineage regulation through the integrating analysis of three high-quality human prostate cancer cohorts (Figure 1C). It is widely accepted that both prostate basal and luminal cells have bi-potential plasticity, which was found in 3 dimensional organoids and UGSM tissue recombination assay (12, 16). In this study, we found that ERG expression strongly facilitates the differentiations towards luminal phenotype in both luminal organoids and basal organoids, consistent with previous findings that ERG expressions induced a significant decrease in the proportion of prostate basal cells (63, 64). Moreover, our current study indicates that luminal cells tend to be more liable for lineage regulation conducted by ERG, when compared with basal cells. Together with their clinical relevance, our findings suggest the important role of ERG in initiation of primary prostate cancer with luminal cell features.

Previous studies have provided some insights into the functional role of androgen receptor (AR) in cell lineage regulation in both normal prostate development and prostate cancer. *In vivo* tissue recombination modeling suggests that stromal AR, but not epithelial AR, is essential for prostate developmental growth and morphogenesis (65, 66). Consistent with these findings, recent mouse lineage-tracing studies have demonstrated that in adult prostate, specific AR deletion in luminal cells has little effects on luminal cell differentiation (50). As for prostate cancer, highly analogous to the previous findings that prostate tumors with AR knockout were characterized by luminal features in mouse models of *Pb-Cre*^*+*^; *Pten*^*flox/flox*^; *AR*^*flox/Y*^ (51) and *Nkx3.1*^*CreERT2/+*^; *AR*^*flox/Y*^; *Pten*^*flox/flox*^; *R26*^*YFP/+*^ (50, 54), our results (Figure 4, A and B) indicate that the luminal lineage differentiations for prostate cancer cells are not directly dependent on AR expression, providing the novel insights that ERG can directly determine the prostate cancer cell luminal lineage through the changes of global chromatin interactions. Consistently, deletions of ERG or ERG specific binding site disrupt the prostate luminal lineage, leading to the differentiation of prostate basal cell lineage (Figure 4, A and B and Figure 6). However, there are also limitations for our study. The percentage of TMPRSS2-ERG fusion is nearly 50% in primary prostate cancer, the other half without ERG expression also display a luminal cell phenotype. Also, both human and mouse normal prostate luminal cells are lack of ERG expression. Therefore, further researches to define other master regulators with the function in ERG-negative prostate cancer or normal prostate lineage regulation are warranted, which may provide rationale for a novel therapeutic strategy and prostate development. Further investigations to dissect cancer-stage-specific roles of luminal-cell AR in both primary prostate cancer and advanced prostate cancer will be really necessary.

Furthermore, PTEN deletion is another common genetic alteration event in primary prostate cancer (31, 67). It was demonstrated that Pten deletion led to basal differentiation, validated by a significant increase of Krt5^+^/Trp63^+^ cells with disease progression (68). Consistently, comparing with wild-type luminal organoids, our Pten null organoids also exhibited basal differentiation (Figure 3A). This could explain the clinical relevance that PTEN loss significantly occurs with ERG fusion, which may facilitate cancer cells to maintain both proliferation capacity and luminal characteristics. Gradually increased incidents of PTEN gene deletion and PI3K signaling pathway activation were identified during prostate cancer progression (48, 49, 69). Therefore, our study demonstrates a possible molecular mechanism underlying the basal lineage plasticity in advanced prostate cancer.

TMPRSS2-ERG translocation represents a distinct subset on the *cis*-regulatory landscape in primary prostate tumors (39). ERG overexpression was known to induce the global changes in chromatin conformation (70). Here, we have further proved that ERG overexpression globally induces chromatin interaction changes (Figure 5, A and B and Supplemental Figure 5, C and D). Moreover, these chromatin interaction changes are associated with the coordinated DEG expressions. Through a distant binding, ERG can regulate Trp63 expression by chromatin interactions. Importantly, deletion of this binding site remarkably reverses the lineage plasticity towards basal differentiation. Compelling data in supporting this hypothesis has also been obtained from the re-analysis on the publicly available human datasets with ERG ChIP-seq, which can validate the conserved existence of ERG binding site in human prostate cells (58). Therefore, we have successfully obtained a novel finding of the conserved ERG binding site that contributes to prostate lineage plasticity. In addition, we have also provided a novel research paradigm for the investigation on how TFs regulate their responsive genes through chromatin interactions instead of direct binding at the gene body regions.

Taken together, ERG is identified as a master transcription factor to manipulate plasticity in prostate cell lineage differentiation towards the pro-luminal programing through chromatin interactions. Our findings can propose a novel working model for elucidating the detailed mechanisms for pursuing a fundamental and long-standing goal aimed at how prostate cancer cells actively maintain luminal lineage identities, as well as for providing the further supporting researches on the role of lineage plasticity in prostate cancer initiation.

## Methods

### Analysis pipeline for identification of the master transcription factors

Prostate cohorts with more than 100 samples and RNA expression profiles were selected for downstream analysis. After prostate cancer cohorts filtering, further cancer subtyping was performed based on the three selected cohorts including FHCRC (158 samples), MSKCC (150 samples) and TCGA (498 samples). For each cohort, integrative classifier was performed to identify the transcription factors which correlated with epigenetics modifications, and PAM50 classifier was performed to define prostate cancer lineage-related transcription factors. In detail, to identify the potential of a TF as master regulator, samples of every cohort were firstly divided into three groups according to this TF expression levels, termed as TF-high, TF-medium and TF-low. Meanwhile, samples were also categorized into another three groups using PAM50 classifier or integrative classifier using their own marker genes respectively. PAM50 classifier was performed originally based on the algorithm(42). We downloaded source code from the University of North Carolina Microarray Database (https://genome.unc.edu/pubsup/breastGEO/). We excluded normal-like subtype and HER2 subtype similarly as previous work(41). Integrative classifier was performed based on 285 genes using unsupervised hierarchical clustering(40). Next, Pearson’s chi-squared test was performed to evaluate the correlation between its expression levels and subtypes, and overlapped TFs in integrative classifier and PAM50 classifier were defined as overlapped TFs. Taking confidence into consideration, overlapped TFs that occurred in at least two cohorts were defined as master TFs for further study. To visualize the significance of each master TF in all of the three cohorts, we performed bubble plot based on transformed p-value of each cohorts 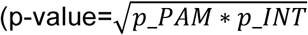, *p_PAM* was calculated by Pearson’s chi-squared test based on PAM50 classifier, *p_INT* was calculated by Pearson’s chi-squared test based on integrative classifier).

### Gene targeting and mouse breeding

All mouse studies are approved by SIBCB Animal Care and Use Committee. Mice were bred and maintained according to Shanghai Laboratory Animal Center Institutional Animal Regulations. *Tmprss2*^*CreERT2/+*^(46), *Pten*^*flox/flox*^(71), *Rosa26*^*EYFP/EYFP*^(72), *Rosa26*^*ERG/ERG*^(38), and *Pb-Cre4*(73) mice were previously described and all in the C57/B6 background. The *Tmprss2*^*CreERT2/+*^; *Rosa26*^*EYFP/EYFP*^ (T2Y), *Tmprss2*^*CreERT2/+*^; *Rosa26*^*EYFP/ERG*^ (T2YE), *Tmprss2*^*CreERT2/+*^; *Pten*^*flox/flox*^; *Rosa26*^*ERG/ERG*^ (T2PE), *Rosa26*^*ERG/ERG*^, *Tmprss2-ERG* knock-in and *Pb-Cre4; Rosa26*^*ERG/ERG*^ mice were generated through standard mouse breeding within the SIBCB animal facility.

### Mouse procedures

For tamoxifen (Toronto Research Chemicals) treatment of T2Y, T2YE and T2PE mice, tamoxifen was dissolved in 20 mg/mL corn oil and injected intraperitoneally to 8-week-old mice at a dose of 3 mg every other day for three doses. Mice were euthanized as indicated timeline after the first tamoxifen dose.

### Isolation and culture of mouse prostate epithelial cells

Mouse prostates were dissected and minced with scissors, then digested with Collagenase/Hyaluronidase (STEMCELL, 07912) for 30 minutes with every 10-minute shaking in cell incubator. And subsequently digested with TrypLE (GIBCO, 12605-028) for another 15 minutes with shaking. DMEM (GIBCO, C11995500BT) with 10% FBS was added to stop the digestion and then cells were centrifuged for collection. Cells were cultured and processed as described previously(12, 74).

### Flow cytometry and fluorescence-activated cell sorting (FACS)

Flow cytometry cell sorting and analysis of mouse prostate cells were performed on an Aria II (BD Biosciences). Single cell suspensions of T2Y mice anterior prostates were stained using CD49f-PE (eBioscience, eBioGoH3, 1:200) and DAPI (4’,6-diamidino-2-phenylindole). *Rosa26*^*ERG/ER*G^ mice prostate cells were stained using a CD49f-PE (eBioscience, eBioGoH3, 1:200), Cd24-FITC (biolegend, M1/69, 1:200) and DAPI. Sorted murine prostate basal cells and luminal cells were fixed in Foxp3 Fixation/Permeabilization buffer (eBioscience, 00-5521-00) and stained with CK5-Alexa Fluor® 488 antibody (Abcam, ab193894, 1:500) or CK8+18-Alexa Fluor® 488 antibody (Abcam, ab192467, 1:500).

### UGSM tissue recombination assays

UGSM tissue recombination assays were performed as described previously(47). Briefly, urogenital sinus mesenchyme (UGSM) cells were dissociated from urogenital sinus of day 18 rat embryos. Added 2-4mL 1% trypsin to the 3.5 cm dish and digested around 20 minutes at 4 degrees until the UGSM was fluffy. Wash the UGSM with DMEM with 10% FBS for twice. Transfer the UGSM cells to 4-6mL 0.1% collagenase B with 1%DNase and incubated at 37 degrees. Shaking vigorously every 10 minutes and carefully collected supernatant, then filtered cell suspension through 70μm strainer. Wash the UGM cells with DMEM with 10% FBS for twice to remove collagenase B. Mix mouse prostate organoids with UGM cells at appropriate ratio. Resuspend the cell mix with collagen for culture overnight. The cell pellets were transplanted under renal capsule of 6-week old SCID mice. The grafts were collected and analyzed as indicated timeline after transplantation.

### Lentiviral CRISPR/Cas9-mediated knock-out

To knock out ERG and AR in mice organoids, we designed three pairs of single guide RNA (sgRNA) sequences for human ERG and mouse AR using the design tool from the Feng Zhang Lab (MIT) and cloned the targeting sequences into the LentiCRISPRv2 vector obtained from Addgene. Lentiviruses for ERG sgRNAs, AR sgRNAs or vector control were generated in 293T cells by standard methods using lentiviral packaging vector. Mice and human prostate cells were infected with lentivirus for 48 hours and selected with 2 μg/mL puromycin for 7 days. ERG protein level and organoids histology were analyzed 21 days after infection. To knock out the distal ERG-binding site with the length of 880 bp, four single guide RNAs were designed including two upstream sgRNAs and two downstream sgRNAs which were subsequently cloned into codon-optimized SpCas9 plasmids PX330-RFP and PM458-GFP (derivative of PX330 backbone) respectively. Transient transfection was conducted to transfect above plasmids into LCD-ERG organoids and *Pten*^*-/-*^; *R26*^*ERG*^ organoids with X-tremeGENE™ 9 transfection reagent (Roche, 6365779001). After 2 days, GFP/RFP double positive cells were sorted into 96-well plate using FACS. Knock-out efficiency was identified by PCR with 3 EB-KO-identify primers and Sanger sequencing. The target guide sequences and EB-KO-identify primers are listed as followed:

sgERG-1-F: CACCGACACCGTTGGGATGAACTA;

sgERG-1-R: AAACTAGTTCATCCCAACGGTGTC;

sgERG-2-F: CACCGTTCCTTCCCATCGATGTTC;

sgERG-2-R: AAACGAACATCGATGGGAAGGAAC;

sgERG-3-F: CACCGTACAGACCATGTGCGGCAG;

sgERG-3-R: AAACCTGCCGCACATGGTCTGTAC;

sgAR-1-F: CACCGGTGGAAAGTAATAGTCGAT;

sgAR-1-R: AAACATCGACTATTACTTTCCACC;

sgAR-2-F: CACCGCACTACGGAGCTCTCACTTG;

sgAR-2-R: AAACCAAGTGAGAGCTCCGTAGTGC;

sgEB-1-F: CACCGATATAGCACCTCGGTTCCCA;

sgEB-1-R: AAACTGGGAACCGAGGTGCTATATC;

sgEB-2-F: CACCGGTGGAAGAGGCATCGAATAG;

sgEB-2-R: AAACCTATTCGATGCCTCTTCCACC;

sgEB-3-F: CACCGATGTGATGCCTTCAGGCACG;

sgEB-3-R: AAACCGTGCCTGAAGGCATCACATC;

sgEB-4-F: CACCGCTGGGAACCGAGGTGCTATA;

sgEB-4-R: AAACTATAGCACCTCGGTTCCCAGC;

EB-KO-identify-F: TTGACAATATTGGAATTAGACGATAT;

EB-KO-identify-R1: AGTCACTCATGAGCAGCGTC;

EB-KO-identify-R2: ACAACAACTTGACCGTGTGG;

sgControl-F: CACCGGGCGAGGAGCTGTTCACCG;

sgControl-R: AAACCGGTGAACAGCTCCTCGCCC;

### Stable gene expression

cDNAs for human prostate cancer *TMPRSS2-ERG* fusion was cloned into retroviral-based vector MSCV-C-HA (Addgene). Retrovirus was produced in 293T cells by standard methods using Ampho packaging vector. Prostate organoids were infected and selected with 2μg/mL puromycin for 7 days at 48 hours after infection for subsequent histology and graft studies.

### Immunohistochemistry

Organoids were fixed in 4% paraformaldehyde (Electron Microscopy Sciences) for 15 minutes at room temperature. Mouse prostates and mouse organoid grafts derived from UGSM tissue recombination assays were fixed using 4% paraformaldehyde overnight at 4 degrees. Organoids and tissues were processed for paraffin embedding using Leica ASP6025 tissue processor (Leica Biosystems). Freshly cut 5 microns paraffin sections were stained on Leica Bond RX (Leica Biosystems) with appropriate negative and positive controls. The following antibodies were diluted in Antibody Diluent (Leica) as indicated: ERG (Abcam, ab92513, 1:100); p63 (Abcam, ab735, 1:500); CK5 (Covance, PRB-160P, 1:2,000); CK8 (Covance, cat# MMS-162P, 1:1,000); PTEN (Cell Signaling Technology, 9188, 1:100); GFP (Abcam, 13970, 1:200); phospho-AKT (Ser473) (Cell Signaling Technology, 4060, 1:50); HA (Cell Signaling Technology, cat# 3724, 1:200); Ki67 (Abcam, cat# ab16667, 1:200).

### Immunofluorescence

Organoids were fixed in 4% paraformaldehyde (Electron Microscopy Sciences) for 15 minutes at room temperature. Mouse prostates were fixed using 4% paraformaldehyde for 2 hours at 4 degrees. Organoids and mouse prostate tissue were embedded using Tissue-Tek O.C.T. Freshly cut 5 Micron paraffin sections were stained with CK5 antibody (Covance, PRB-160P, 1:1,000); CK8 antibody (Covance, cat# MMS-162P, 1:1,000); ERG (Abcam, ab92513, 1:100); p63 (Abcam, ab735, 1:500) on Leica Bond RX (Leica Biosystems) with appropriate negative and positive controls. After washing in PBS, slides were mounted with Mowiol® 4-88 (Millipore, 475904) and imaged with a Leica TCS SP5 II confocal microscope. Immunofluorescence was independently performed twice.

### Target cell quantification

To calculate the number of Trp63 and ERG, Krt5 and ERG double positive cells in the T2PE mice prostate PIN lesions and intraductal carcinomas, slides were scanned with Pannoramic Confocal Scanner (3DHistech, Hungary) using 40x/1.2 water objective. Appropriate areas of the scanned tissue were exported to .tiff files and analyzed in ImageJ/FIJI (NIH). Appropriate thresholds were applied for each channel, and cells expressing ERG were segmented out from the T2PE mice prostate PIN lesions and intraductal carcinomas. Then analysis was performed to determine the percentage of those cells were also positive for basal markers Trp63 or Krt5.

### Western blotting

Cell lysates were prepared in RIPA buffer supplemented with proteinase/phosphatase inhibitor. Proteins were resolved in NuPAGE Novex 4–12% Bis-Tris Protein Gels (Life Technologies) and transferred electrophoretically onto a PVDF 0.45 mm membrane (Millipore). Blocking was conducted for 1 hour at room temperature in 5% milk in TBST buffer and were incubated overnight at 4 degrees with the primary antibodies diluted in 5% milk in TBST buffer. The following primary antibodies were used: β-Actin (Sigma-Aldrich, A3854, 1:5,000), ERG (Abcam, ab92513, 1:1,000); CK5 (Covance, PRB-160P, 1:1,000); CK8 (Covance, MMS-162P, 1:1,000); PTEN (Cell Signaling Technology, 9188, 1:1,1000); phospho-AKT (Ser473) (Cell Signaling Technology, 4060, 1:1,1000). Immunoblots were independently performed at least twice.

### Quantitative RT-PCR analysis

Total RNA was extracted with TRIzol reagent (ambion, 15596018) and reverse transcription was further performed with 500 ng total RNA as initiation material with PrimeScriptTM RT Master Mix (TaKaRa, RR036A). qRT-PCR was conducted with 2 x S6 Universal SYBR qPCR Mix (NovaBio, Q204) using the manufacture’s protocol. The primers sequences are listed as followed:

Krt5-qPCR-F: GAACAGAGGCTGAGTCCTGGTA;

Krt5-qPCR-R: TCTCAGCCTCTGGATCATTCGG;

Krt14-qPCR-F: GAAGAACCGCAAGGATGCTGAG;

Krt14-qPCR-R: TGCAGCTCGATCTCCAGGTTCT;

Trp63-qPCR-F: GTATCGGACAGCGCAAAGAACG;

Trp63-qPCR-R: CTGGTAGGTACAGCAGCTCATC;

Krt8-qPCR-F: TGGAAGGACTGACCGACGAGAT;

Krt8-qPCR-R: GGCACGAACTTCAGCGATGATG;

Krt18-qPCR-F: AATCAGGGACGCTGAGACCACA;

Krt18-qPCR-R: GCTCCATCTGTGCCTTGTATCG.

### RNA-seq data processing and analysis

RNA sequencing libraries were prepared with the VAHTS mRNA-seq V3 Library Prep Kit for Illumina® (Vazyme, NR611). Sequencing was performed by Berry Genomics. Low-quality sequences and adapters were filtered by cutadapt-1.15. Clean reads were mapped to the mm9 genome using hisat2-2.1.0(75). Gene expression was quantified at the gene level using featureCounts(76). Differentially expressed genes (DEGs) were analyzed by DESeq2(77) using raw counts. And adj.P.value < 0.05 was set as threshold to define DEGs. GSEA (Gene Set Enrichment Analysis)(78) was conducted to determine statistically significant defined signatures based on the normalized expression value calculated from DESeq2. Enriched pathway analysis was performed using Metascape(79).

### ATAC-seq library preparation

50,000 cells were collected by centrifuging and washed with ice-cold PBS. Cells were lysed in 50μL ice-cold lysis buffer (10mM pH7.4 Tris-HCl; 10mM NaCl; 3mM MgCl2; 0.5%NP-40) for 10 minutes on ice. Immediately after lysis, nuclei were spun at 500g for 5 minutes using a refrigerated centrifuge at 4 degrees. Following steps to generate sequencing libraries was performed with TruePrep DNA Library Prep Kit V2 for Illumina (Vazyme, TD501).

### ATAC-seq data processing and analysis

We used Partek Genomics Suite to map sequencing reads and remove duplicate reads to mouse reference genome mm9. Peaks were identified using HOTSPOT with default parameters (http://www.uwencode.org/proj/hotspot/). HOTSPOT analysis generates two types of peaks: narrow peak and hotspot regions (broad peak). In this study, we used the narrow peaks for downstream analysis.

### Quantify chromatin accessibility of ATAC-seq

We referred to narrow peak as regulatory element (RE). ATAC-seq will measure the accessibility in a given regulatory element. We can quantify the openness for the RE by a simple fold change score, which computes the enrichment of read counts by comparing the RE with a large background region. Briefly, let N be the number of reads in RE of length L and G that in the W background window (1Mb in our case) around this RE. The openness of RE o can be defined as: 

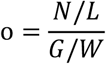

### Differential ATAC-seq peaks analysis

Differential ATAC-seq peaks analysis was performed by comparing LCD organoids with LCD-ERG organoids, as well as comparing *Pten*^*-/-*^ organoids with *Pten*^*-/-*^; *R26*^*ERG*^ organoids. We defined the sample specific peaks with >1.5 fold-changes and an openness value >1.

### ChIP-seq library preparation

10 million target cells were collected for centrifuging and then resuspended in 10mL freshly made 1% formaldehyde with incubation at room temperature for 10 minutes with rotation. 526ul 2.5M glycine was added to a final concentration of 125 mM to quench the formaldehyde at room temperature for 5 minutes with rotation. Cells were pelleted and washed in ice-cold PBS. 880μL of ice-cold cell lysis buffer (1% SDS; 10 mM EDTA; 50 mM Tris-HCl; 1X proteinase inhibitor) was added to lyse cross-linked cells with rotation at 4 degrees for 30 minutes. 880μL cell lysis was transferred into a Covaris milliTUBE 1mL AFA Fiber and sheared with Covaris S220 (Fill level: 10; Duty cycle: 5; PIP: 140; Cycles/Burst: 200; Time: 4 minutes). Samples was clarified for 15 minutes at 16100 rcf at 4 degrees. Another 1600μL ChIP Dilution Buffer (0.01% SDS; 1.1% Triton X-100 (fisher scientific, BR151-500); 1.2 mM EDTA; 16.7 mM Tris pH 7.5; 167 mM NaCl) was added to achieve SDS concentration of 0.33%. 60μL Protein A beads was pre-cleared in 500μL ChIP Dilution Buffer. Protein A beads (Invitrogen, 10001D) were resuspended in 60μL of ChIP Dilution Buffer and added to sample at 4 degrees for 1 hour. Samples with beads was put on magnet and the supernatant was transferred into new tubes. Target antibody (5 μg for H3K27ac (Abcam, ab4729), 8 μg for ERG (Abcam, ab92513)) was added at 4 degrees for incubation overnight with rotation. To bind target antibody, 60μL pre-cleared Protein A beads were added to samples with two-hour rotation at 4 degrees. Beads was washed three times each with Low Salt Wash Buffer, High Salt Wash Buffer and LiCl Wash Buffer. 100μL of freshly-made DNA Elution Buffer (50 mM NaHCO_3_; 1% SDS) was added to resuspend ChIP sample beads with incubation at RT for 10 minutes, followed by 3 minutes at 37 degrees. ChIP sample beads were placed on magnet and the supernatant was transferred to a new tube. Another 100μL of DNA Elution Buffer was added to ChIP samples and the same incubation protocol was conducted. Supernatant of ChIP samples was collected again to the new tube. 10μL of Proteinase K (Invitrogen, 25530049) was added to each sample with incubation at 67 degrees at least 4 hours with shaking. DNA was purified with QIAGEN purification kit (QIAGEN, 28106) and eluted in 20μL of nuclease-free water. Sequencing libraries were prepared from TruePrep DNA Library Prep Kit V2 for Illumina (Vazyme, TD503) using the manufacturer’s recommended protocol.

### ChIP-seq data processing and analysis

The ChIP-seq pipeline was based on the ENCODE (phase-3) transcription factor and histone ChIP-seq pipeline specifications (by Anshul Kundaje) (https://github.com/ENCODE-DCC/chip-seq-pipeline2). Sequencing reads were mapped to mm9 using bwa(80) with parameters *aln -q 5 -l 32 -k 2*. Unmapped, multimapping, low-quality reads and duplicates were filtered by samtools(81). Peaks were identified using MACS2(82) with an FDR of 0.05.

### Bridge Linker-Hi-C (BL-Hi-C) assay

The libraries of BL-Hi-C were generated using the two-step ligation protocols as previously described(55). One million cells were cross-linked in 1mL 1% methanol free formaldehyde (Sigma-Aldrich, F8775) with shaking for 10 minutes at room temperature. 2.5M glycine was added to quench formaldehyde to a final concentration of 0.2M with shaking for 10 minutes at room temperature, and then the cells were put on ice for 5 minutes. Cell pellets were collected with centrifuging and then 1mL 0.1% SDS BL-Hi-C Lysis buffer (50 mM HEPES-KOH, pH 7.5; 150 mM NaCl; 1 mM EDTA; 1% Triton X-100; 0.1% Sodium Deoxycholate; 0.1% SDS) with proteinase inhibitor was added with incubation for 15 minutes at 4°C with shaking at 850 rpm. Cells were centrifuged and resuspended with 1mL of 0.55% SDS BL-Hi-C Lysis buffer (50 mM HEPES-KOH, pH 7.5; 150 mM NaCl; 1 mM EDTA; 1% Triton X-100; 0.1% Sodium Deoxycholate; 1% SDS; 1X proteinase inhibitor) for another 15-minute incubation at 4°C with shaking at 850 rpm. Cells were collected by centrifuging and resuspended with 100 µL of 0.3% SDS in 1×NEBuffer 2.1(NEB, B7002S) with following shaking at 37°C for 30 minutes at 900 rpm. 135µL nuclease-free water and 15μL 20% Triton X-100 were added and samples were kept at 37°C for 15min with shaking at 900rpm. Cell pellets were collected and resuspended by 76.5μL nuclease-free water, 10μL 10 X NEBuffer2.1, 2.5μL of 20% Triton X-100, 1μL BSA (100x, 10mg/mL) and 10μL HaeIII (10U/μL, NEB, R0108S) with shaking at 37°C overnight. 2μL 10mM dATP (NEB, N0440S) and 5μL Klenow Fragment (3′->5′ exo-, NEB, N0202S) for A-tailing were added at 37°C for 40 minutes. Cell pellets were collected and resuspended with 411μL nuclease-free water, 50μL T4 DNA ligase buffer (NEB, B0202S), 25μL 20% Triton X-100, 5μL BSA, 5μL T4 DNA ligase (NEB, M0202S) and 4μL Bridge Linker (Bridge Linker S2-F: /5P/CGCGATATC/iBIOdT/TATCTGACT; Bridge Linker S2-R: /5P/GTCAGATAAGATATCGCGT) with subsequent rotation at room temperature for 4 hours. Cell pellets were centrifuged at 1000g for 2 minutes at 4°C and resuspended with 88μL nuclease-free water, 10μL λ Exonuclease Buffer, 1μL λ Exonuclease (NEB, M0262L) and 1μL Exonuclease I (NEB, M0293L) with shaking at 37°C for 60 minutes. DNA purification was conducted with Phenol: Chloroform: Isoamyl Alcohol 25:24:1(Sigma-Aldrich, cat. no. P3803) and eluted with 60μL of TE buffer. Add another 60μL 2XB&W buffer was added into DNA solution for sonication with setting Covaris parameters for the DNA size of 300 bp. Streptavidin C1 beads were washed (Invitrogen, 65001) with 1mL 1X TWB buffer for twice and resuspended with 20μL 1× B&W buffer. To bind target DNA, beads suspension was added into 130μL sonicated DNA solution with incubation at room temperature for 15 minutes with rotation at 900 rpm. Beads was then washed with 1X TWB buffer, BW buffer and nuclease-free water. Finally, DNAs-on-beads was resuspended in 50μL nuclease-free water. Library construction for sequencing was conducted with VAHTS Universal DNA Library Prep Kit for Illumina (Vazyme, ND607). The BL-Hi-C library was sequenced with the Illumina Sequencer NovaSeq (PE 2×150 bp reads).

### BL-Hi-C data processing

We first trimmed the linkers of BL-Hi-C sequence using trimLinker function of ChIA-PET2(83). HiC-Pro(84) was then performed to process Hi-C data through several main steps including mapping raw reads to mm9 reference genome, detecting valid ligation products, quality control and generating raw contact maps. To evaluate the library quality, we firstly removed duplicated reads, and then divided the remaining valid reads into several groups including cis long-range (>200k), cis short-range (<200k), and trans contacts, each group was indicated by different color. High quality of our BL-Hi-C libraries was demonstrated by ∼40% for the proportion of long-range valid pairs (84). For raw contact matrix generation, in detail, we set a variety of resolutions including 10kb, 20kb, 40kb, 150kb, 500kb, and 1Mb, which denoted that genome was divided into bins with the above equal sizes respectively. Then we used the iterative correction method to eliminate systematic basis to get the normalized contact matrix. Homer (85) was performed to further identify significant interactions (loops) (FDR <0.0001) based on contact maps using 10k resolution.

### Identification of differential chromatin interactions

We performed differential chromatin interactions by comparing LCD organoids with LCD-ERG organoids, as well as by comparing *Pten*^*-/-*^ organoids with *Pten*^*-/-*^; *R26*^*ERG*^ organoids. We referred to the set of significant interactions identified by Homer in each sample as its loop set and differential loops were defined by the difference of two sets.

### Identification of the relationship between chromatin interactions and ERG binding

To characterize the relationship between chromatin interaction and ERG binding, we divided the genome into 1-Mb bins and sorted the genomic bins by the number of *Pten*^*-/-*^; *R26*^*ERG*^/LCD-ERG-specific loops. We next performed Pearson correlation based on the number of *Pten*^*-/-*^; *R26*^*ERG*^/LCD-ERG-specific loops and ERG binding density, generated from BL-Hi-C and ERG ChIP-seq data respectively. We also calculated the average number of ERG binding sites for the first N bins with *Pten*^*-/-*^; *R26*^*ERG*^/LCD-ERG-specific loops (N ranges 20 to 2,779). Results showed a strong association between hotspots of differential chromatin interactions and enrichment of ERG binding.

### Statistical analysis

For significant tests, two-tail Student’s t-test and Pearson’s Chi-squared test were used for comparing differences between two groups, and one-way ANOVA tests were used for multiple groups.

## Acknowledgements

We would like to acknowledge Baojin Wu, Guoyuan Chen and Wei Tang for the animal husbandry and Wei Bian for technical help at the SIBCB Core Facility. We would like to thank the Genome Tagging Project (GTP) Center, Shanghai Institute of Biochemistry and Cell Biology, CAS for technical support. We thank the MSKCC Molecular Cytology (Ning Fan, Mesruh Turkekul, Sho Fujisawa), MSKCC Integrated Genomics Operation (Daoqi You, Agnes Viales) and MSKCC Epigenomics Core Facility (Yang Li). We thank Dr. Wilbert Zwart and Dr. Suzan Stelloo at the Netherlands Cancer Institute for sharing analysis methods of the integrative classifier. This study was supported by grants from the Strategic Priority Research Program of the Chinese Academy of Sciences (XDB19000000 and XDA16020905), the National key research and development program of China (No. 2017YFA0505500), the National Natural Science Foundation of China (81830054 and 81772723), and the US National Cancer Institute (R01CA208100, R01CA193837, P50CA092629 and P30CA008748).

## Author contributions

D.G. and Y.C. conceived and designed the experimental approach. Y.W., B.C. and P.C. provided advice about experimental design. F.L., W.D. and X.Y.X. performed most of the experiments. Q.Y.Y., F.L., L.L. and Y.W. contributed to the computational analysis and statistical analysis. C.F.L. and J.H. generated the expression vectors and lentiviral CRISPR/Cas9 vectors. Z.L., N.H.M., Y.G.L., W.X.G. and S.Q.W. prepared mouse organoids RNA and sequencing. X.Y.Z., Z.L., Y.Q.Z. and R.A. helped with allograft experiments and mouse experiments, D.G. F.L. and Y.C. wrote the manuscript. All authors discussed result and edited the manuscript.

## Accession numbers

The raw data for RNA-seq, ChIP–seq, BL-Hi-C and processed data for ATAC-seq have been deposited in NODE (http://www.biosino.org/node). All data can be viewed in by pasting the accession (OEP000693) into the text search box or through the URL: http://www.biosino.org/node/project/detail/OEP000693, including ATAC-seq data (OEX002111), ChIP-seq data (OEX002110), RNA-seq data (OEX002109) and BL-Hi-C data (OEX002216).

## Supplemental material

Supplemental Table 1. Overlapped TFs for each prostate cancer cohorts.

Supplemental Table 2. 154 Master TFs.

Supplemental Table 3. ERG-upregulating luminal signature genes and ERG-downregulating basal signature genes.

