## Supplementary Figures for "ERG orchestrates chromatin interactions to drive prostate cell fate reprogramming"

### Supplemental Figure. 1

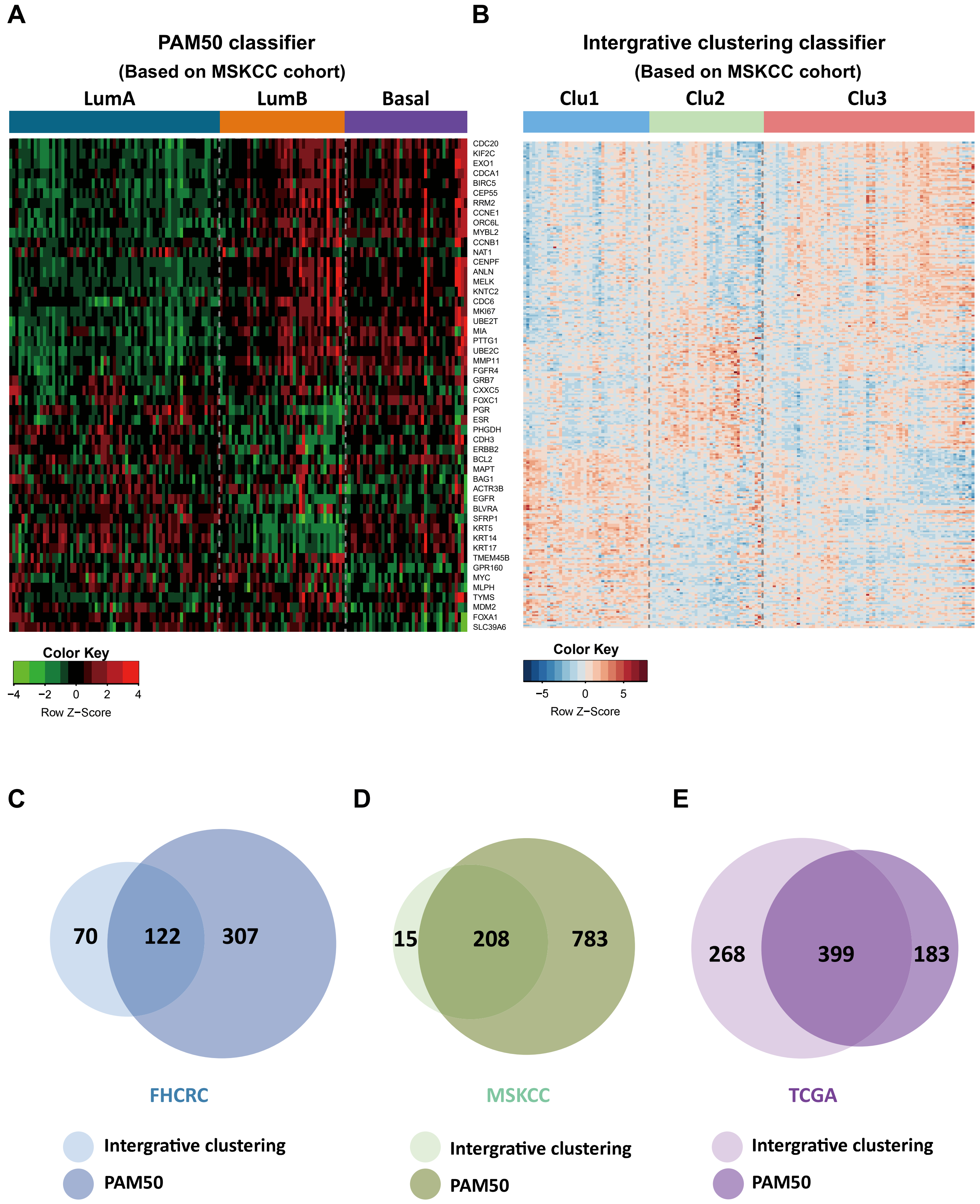

**Supplemental Figure 1 (related to Figure 1). Identification for the master transcription factors with the combination analysis of integrative classifier and PAM50 classifier. (A)** Heatmap showing the expression of PAM50 classification marker genes in each sample of MSKCC cohort, with the results of three subtypes including Luminal A (dark blue), Luminal B (orange) and Basal (purple). The genes are represented with the same order as shown in (41). **(B)** Heatmap showing the expression of integrative classification marker genes in each sample of MSKCC cohort, with the results of three subtypes including Clu 1 (light blue), Clu 2 (light green) and Clu3 (red). The genes are represented with the same order as shown in (40). **(C-E)** Venn diagram showing the number of the overlapped TFs identified by integrative classifier and PAM50 classifier in three cohorts: 122 overlapped TFs of FHCRC cohort **(C)**, 208 overlapped TFs of MSKCC cohort **(D)**, 399 overlapped TFs of TCGA cohort **(E)**.

### Supplemental Figure. 2

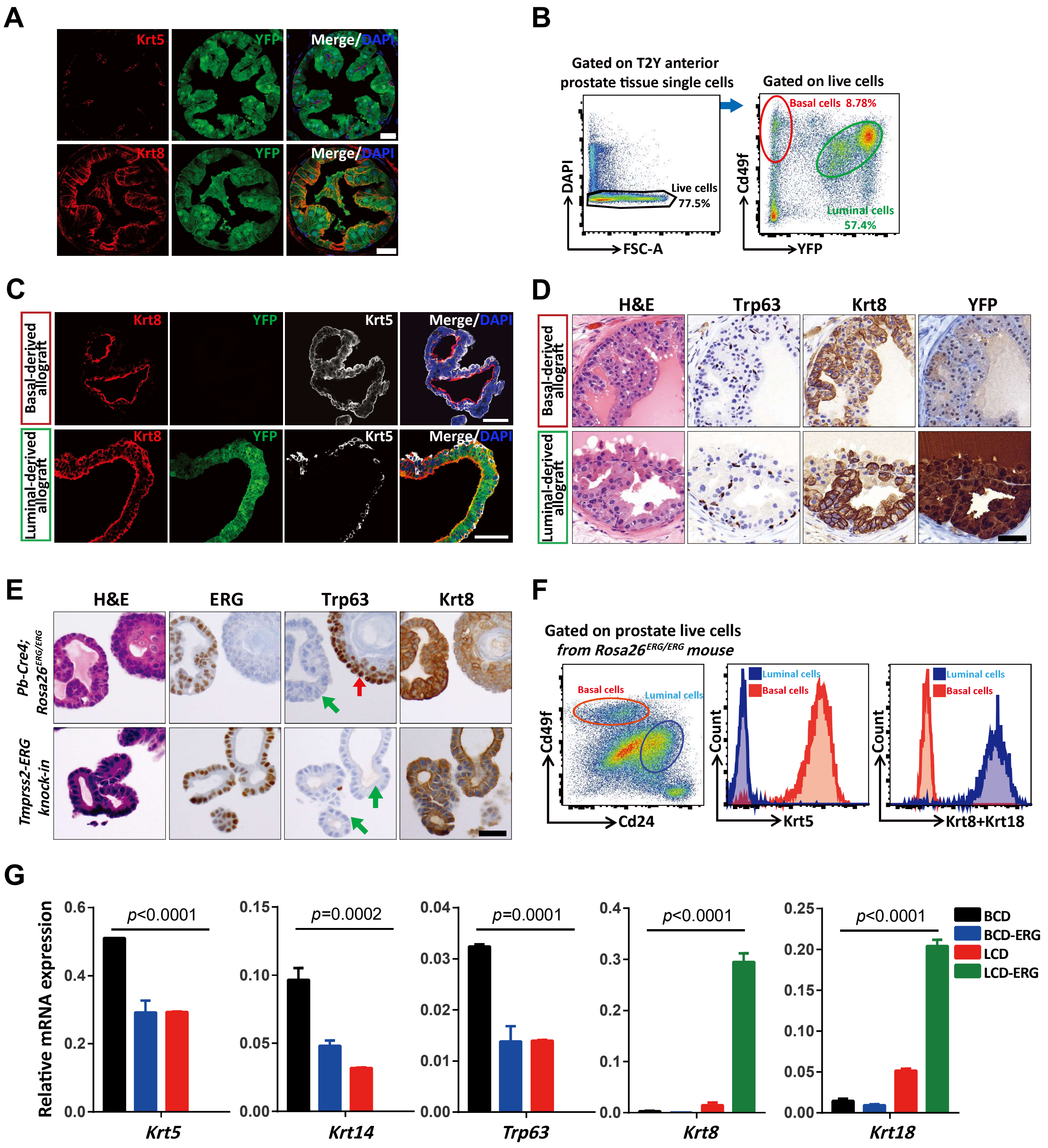

**Supplemental Figure 2 (related to Figure 2). Normal prostate epithelial cells have multiple cell fate and ERG overexpression promotes prostate luminal lineage differentiation. (A)** Immunofluorescence staining of Krt5, Krt8 and YFP in T2Y mice prostates. **(B)** Isolation of $Cd49f^{low}/YFP^{+}$ luminal cells and $Cd49f^{high}/YFP^{-}$ basal cells by flow cytometry. **(C)** Immunofluorescence staining of Krt8, YFP and Krt5 of basal-cell-derived organoids (top) and luminal-cell-derived organoids (bottom). **(D)** H&E and Trp63, Krt8 and YFP IHC staining of allografts from UGSM tissue recombination assays after transplantation of basal- (top) or luminal-cell-derived organoid (bottom) generated from T2Y mice anterior prostate. **(E)** H&E and ERG, Trp63 and Krt8 IHC staining of prostate organoids derived from *Pb-Cre4; Rosa26*$^{ERG/ERG}$ and *Tmprss2-ERG* knock-in mice, respectively. Trp63-negative organoids with ERG expression were indicated by green arrows and Trp63-positive organoids without ERG expression were indicated by red arrows. **(F)** Sorting strategy for prostate basal ($Cd49f^{high}/Cd24^{low}$) and luminal ($Cd49f^{low}/Cd24^{high}$) cells from *Rosa26*$^{ERG/ERG}$ mice (left), intracellular flow cytometry for basal cell lineage marker Krt5 (middle) and luminal cell lineage markers Krt8/Krt18 (right) on *Rosa26*$^{ERG/ERG}$ mice prostate cells. **(G)** QRT-PCR analysis for mRNA expression of basal cell lineage markers *Krt5*, *Krt14*, *Tr63* and luminal cell lineage markers *Krt8*, *Krt18* (one-way ANOVA, mean ± sem). Scale bars, 50 μm.

#### Supplemental Figure. 3

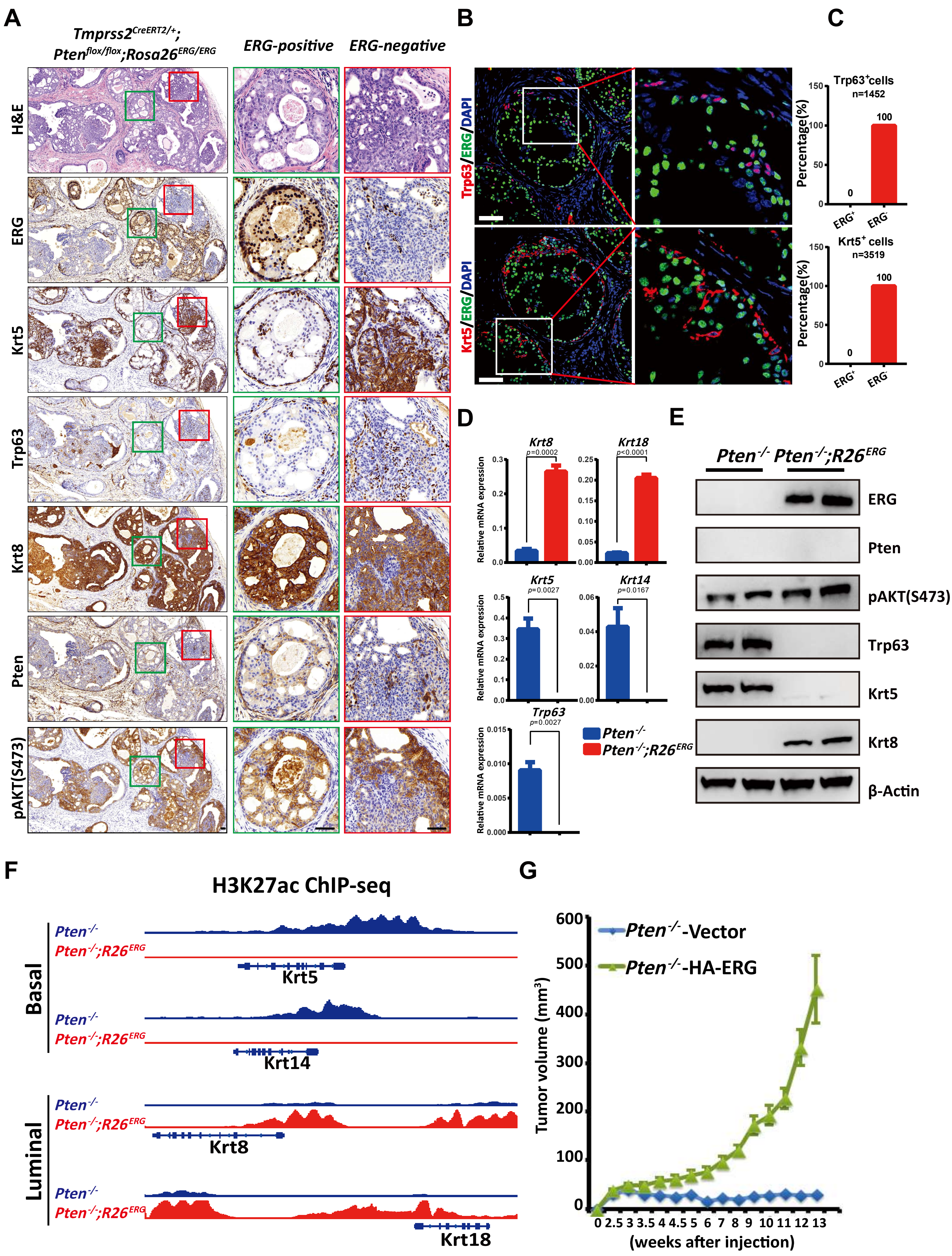

**Supplemental Figure 3 (related to Figure 3). ERG regulates prostate cancer cell lineage in the context of Pten loss. (A)** H&E and ERG, Krt5, Trp63, Krt8, Pten and pAKT(S473) IHC staining of T2PE mice prostate tumors with 7 months post tamoxifen injection, green box indicates ERG-positive region and red box indicates ERG-negative region. **(B)** Immunofluorescence staining of ERG, Krt5 and Trp63 in T2PE mice prostate with 7 months post tamoxifen injection. **(C)** Histogram statistic for the percentage of $ERG^{+}$ cells and $ERG^{-}$ cells in $Trp63^{+}$ cells (top) and $Krt5^{+}$ cells (bottom) respectively. **(D)** QRT-PCR analysis for mRNA expression of basal cell lineage markers *Krt5*, *Krt14*, *Trp63* and luminal cell lineage markers *Krt8*, *Krt18* (two-tailed t-test, mean ± sem). **(E)** Western blotting analysis of ERG, Pten, pAKT(S473), Trp63 and Krt5 in $Pten^{-/-}$ and $Pten^{-/-}$; $R26^{ERG}$ organoids. **(F)** Snapshot representation of the H3K27ac peak profile (defined by H3K27ac ChIP-seq) associated with prostate basal cell lineage markers Krt5, Krt14 and luminal cell lineage markers Krt8, Krt18. **(G)** Subcutaneous tumor growth in SCID mice of Pten loss organoids overexpressing TMPRSS2-ERG fusion protein with HA tag or a control vector respectively. Error bars, mean ± sem, n=10 tumors. Scale bars, 50 μm.

#### Supplemental Figure. 4

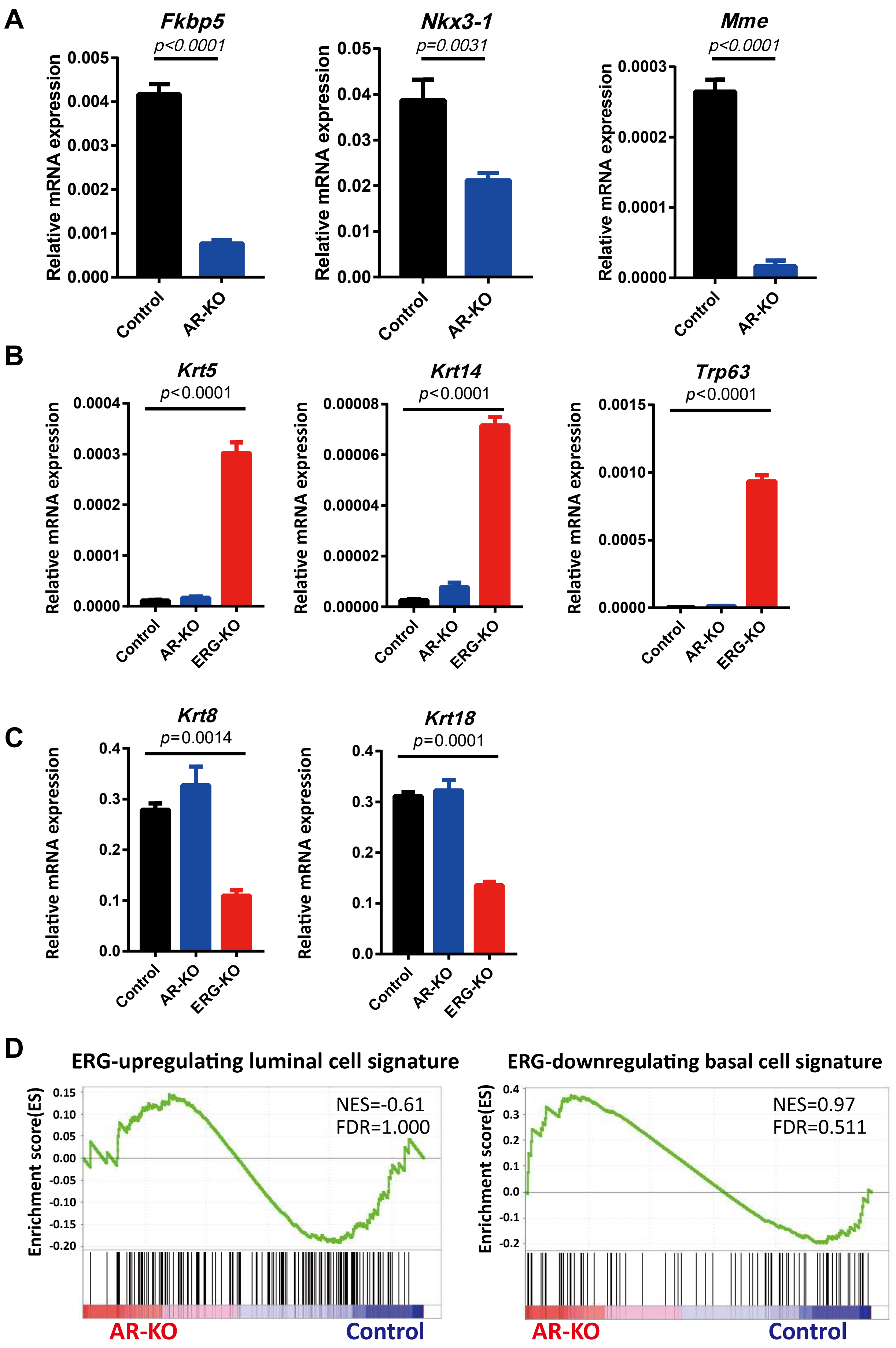

**Supplemental Figure 4 (related to Figure 4). ERG instead of AR is critical for prostate cancer cell lineage regulation in Pten loss context. (A)** QRT-PCR analysis for mRNA expression of AR target genes *Fkbp5*, *Nkx3-1* and *Mme* (two-tailed t-test, mean ± sem). **(B-C)** QRT-PCR analysis for mRNA expression of basal cell markers *Krt5*, *Krt14* **(B)** and *Trp63* and luminal cell markers *Krt8*, *Krt18* **(C)** (one-way ANOVA, mean ± sem). **(D)** GSEA enrichment plot of AR-KO versus Control using ERG-upregulating luminal cell signature genes (left) and ERG-downregulating basal cell signature genes (right) respectively.

#### Supplemental Figure. 5

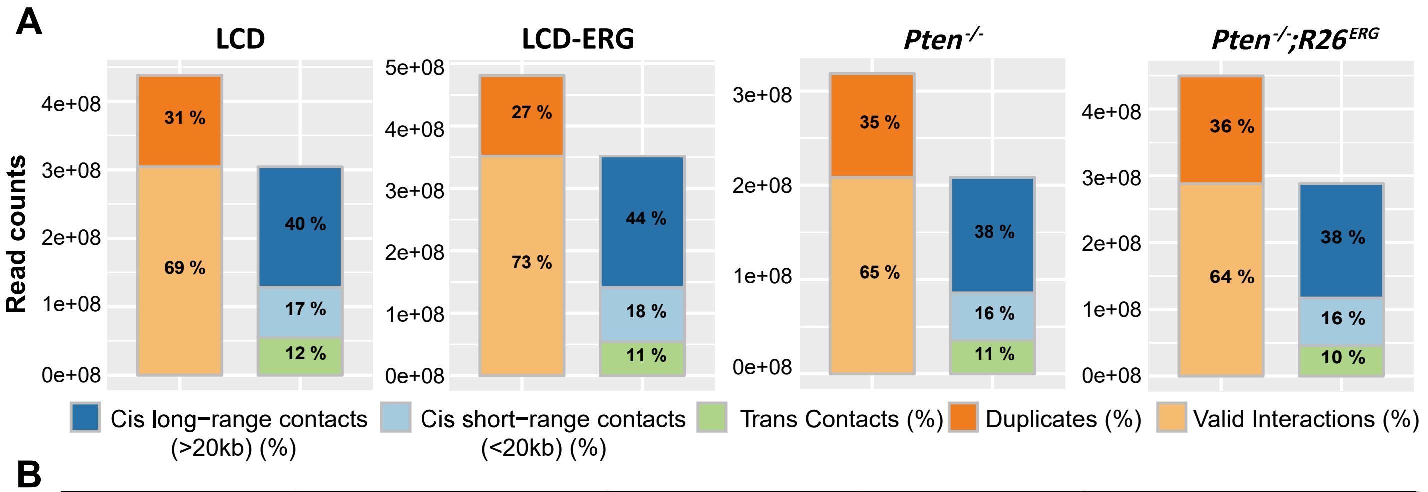


B

| Sample | Unique Pairs | Valid Pairs | Cis Pairs | Percentage of Cis |
|---|---|---|---|---|
| LCD | 487,636,911 | 304,119,352 | 249,087,773 | 81.90% |
| LCD-ERG | 517,797,373 | 351,929,491 | 297,643,740 | 84.57% |
| *Pten*$^{-/-}$ | 380,991,969 | 208,194,264 | 172,818,759 | 83.01% |
| *Pten*$^{-/-}$;*R26*$^{ERG}$ | 520,754,171 | 288,278,089 | 242,728,002 | 84.20% |

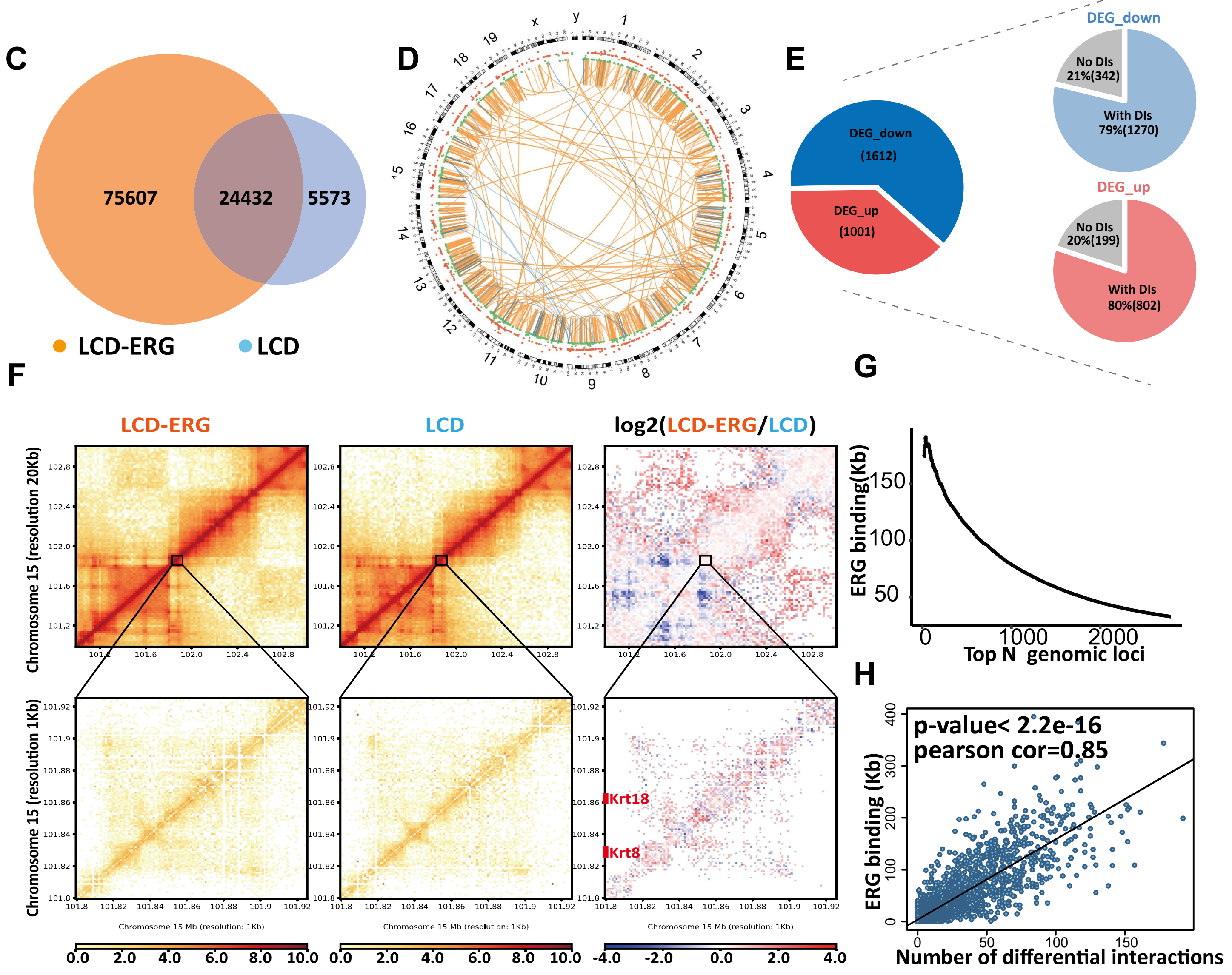

**Supplemental Figure 5 (related to Figure 5). ERG alters chromatin interactions associated with gene expression in normal prostate epithelial cells. (A)** Histogram statistics for the composition of valid pairs in the libraries of LCD, LCD-ERG, *Pten*$^{-/-}$ and *Pten*$^{-/-}$*; R26*$^{ERG}$ organoids respectively. **(B)** Statistics on the number of unique pairs, valid read pairs, cis pairs and percentage of cis pairs in BL-Hi-C libraries of LCD, LCD-ERG, *Pten*$^{-/-}$ and *Pten*$^{-/-}$*; R26*$^{ERG}$ organoids respectively. **(C)** Venn plot showing differential chromatin interactions between LCD and LCD-ERG organoids. **(D)** Circos plot depicting chromosomes 1-19, X and Y on the basis of BL-Hi-C data and RNA-seq data, indicating DIs including LCD-ERG-specific DIs(orange) and LCD-specific DIs(light blue), DEGs including up-regulated DEGs of LCD-ERG (red) and down-regulated DEGs of LCD-ERG (green), respectively. **(E)** Venn plot showing the percentage of down-regulated DEGs(top) and up-regulated DEGs with DIs(bottom). **(F)** The normalized interaction heatmaps of LCD-ERG (left), LCD (middle), and the difference (right) at 20 kb resolution (top) and 1 kb resolution (bottom) of chromosome 15, including Krt8 and Krt18 genomic region. **(G)** Plot showing the density of ERG binding (kb) at each of the ranked (N) differential interacting chromatin loci of 1-Mb intervals. **(H)** Correlation plot showing the positive relationship between ERG binding density and the number of DIs in 1-Mb intervals.

### Supplemental Figure. 6

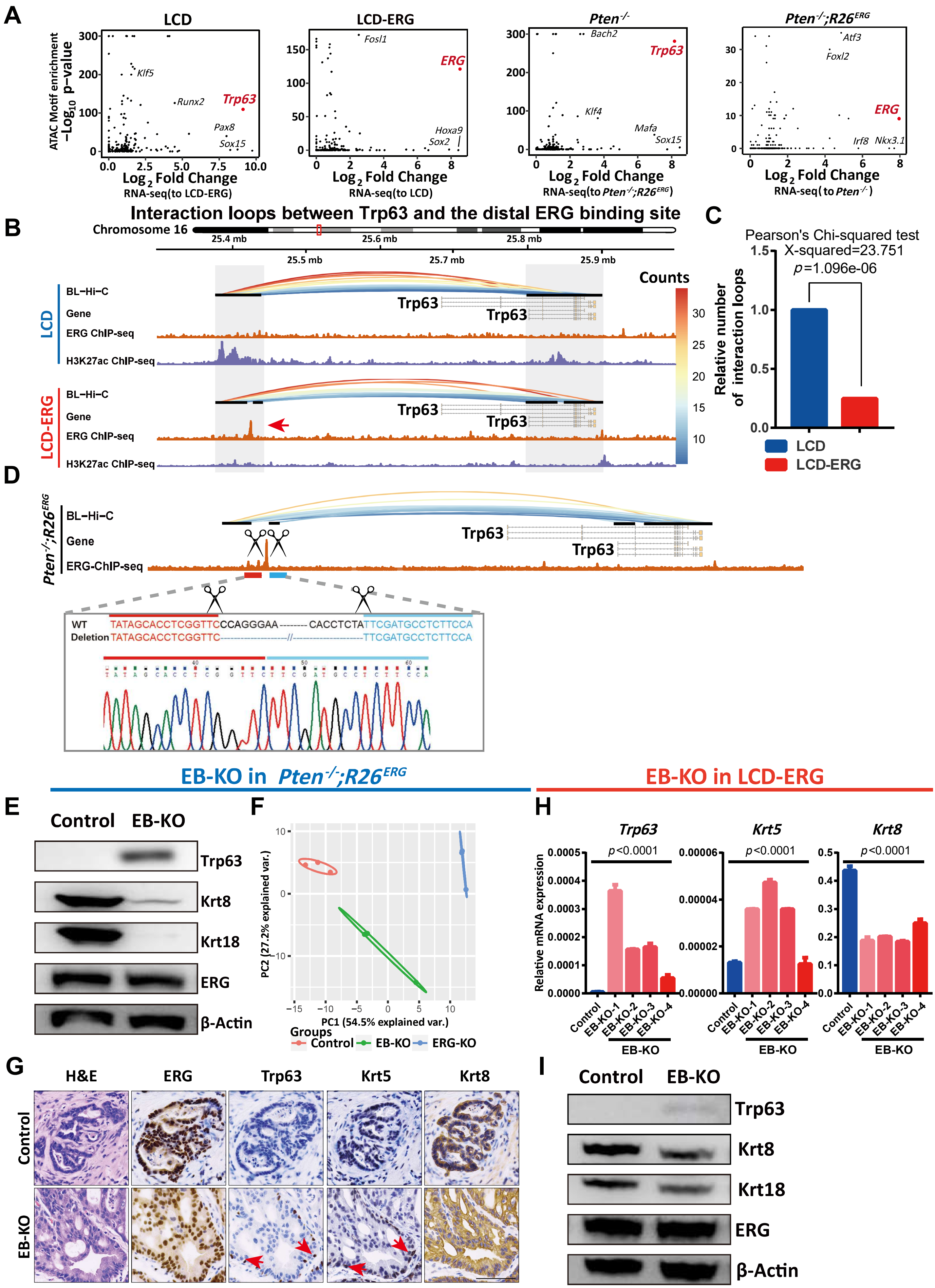

**Supplemental Figure 6 (related to Figure 6). Integrating analysis of ATAC-seq and RNA-seq reveals Trp63 and ERG as master regulators and ERG binding site knock-out impairs luminal lineage phenotype. (A)** Dot plots for transcription factors showing the p-value of motif enrichment analysis and mRNA expression changes from ATAC-seq data and RNA-seq data respectively in LCD, LCD-ERG, *Pten*$^{-/-}$ and *Pten*$^{-/-}$*; R26*$^{ERG}$ organoids respectively. **(B)** 3D signal of BL-Hi-C showing chromatin interactions between the distal ERG binding site and Trp63 gene body region in LCD (top) and LCD-ERG (bottom) organoids respectively. Red arrow indicates the distal ERG binding site. **(C)** Pearson's Chi-squared test to evaluate the differences of interaction loops density between LCD and LCD-ERG organoids. **(D)** Schematic diagram for the strategy to delete the distal ERG-binding site(top) and Sanger sequencing for identification of knock-out efficiency (bottom). **(E)** Western blotting analysis of Trp63, Krt8, Krt18 and ERG in EB-KO and Control of *Pten*$^{-/-}$*; R26*$^{ERG}$ organoids. **(F)** PCA plot for EB-KO, ERG-KO and Control organoids using prostate cell lineage signature genes. **(G)** H&E and ERG, Trp63, Krt5, Krt8 IHC staining of allografts derived from EB-KO and Control of *Pten*$^{-/-}$*; R26*$^{ERG}$ organoids using UGSM tissue recombination assays. **(H)** QRT-PCR analysis of mRNA expression of *Trp63*, *Krt5* and *Krt8* in EB-KO and Control of LCD-ERG organoids (one-way ANOVA, mean ± sem). **(I)** Western blotting analysis of Trp63, Krt8, Krt18 and ERG in EB-KO and Control of LCD-ERG organoids. Scale bars, 50 μm.

#### Supplemental Figure. 7

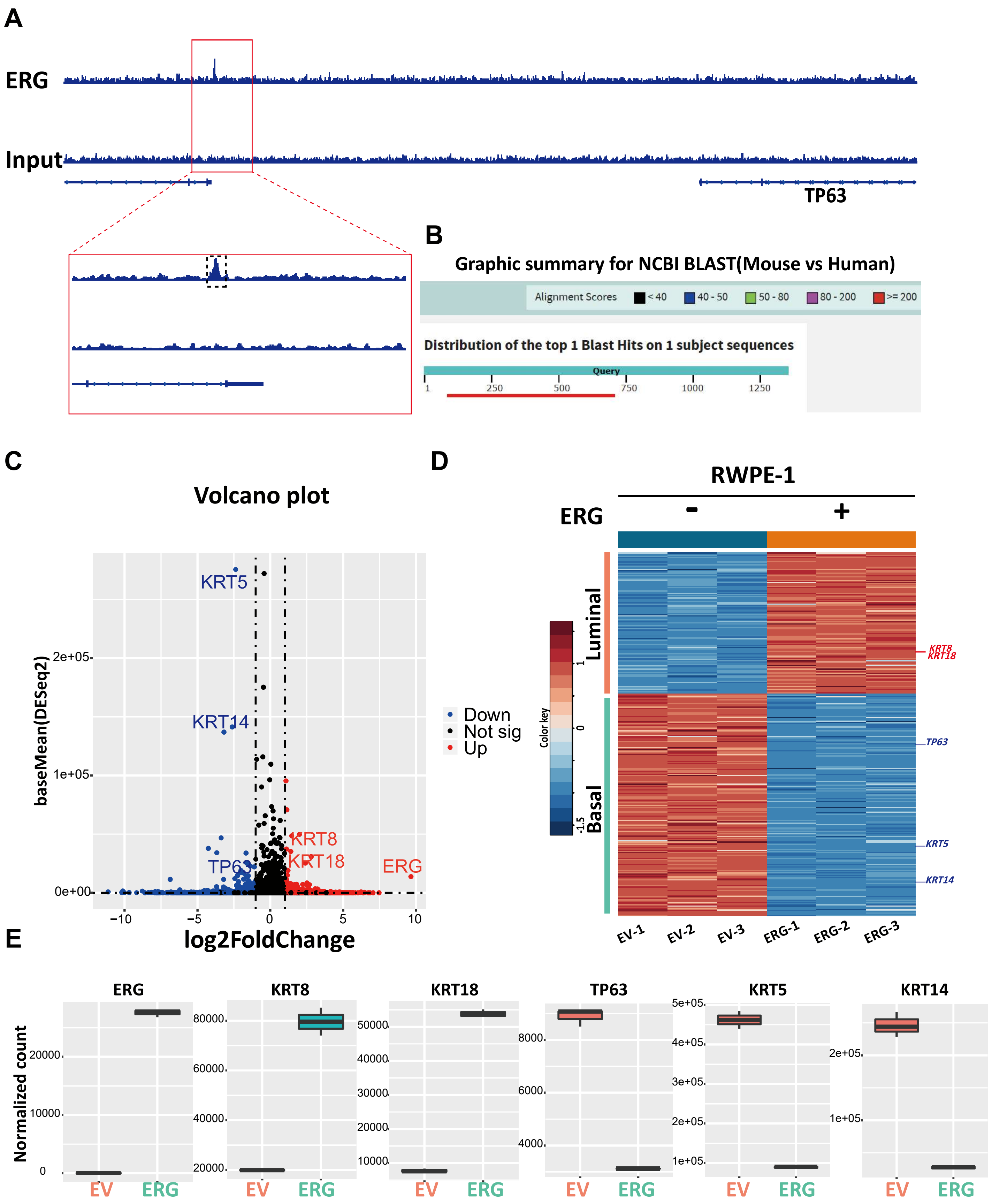

**Supplemental Figure 7 (related to Figure 6). Identification of the conserved distal ERG binding site in human prostate cells. (A)** Snapshot representation of the ERG binding peak profile (defined by ERG ChIP-seq) in RWPE-1 cells with ERG expression. **(B)** Graphic summary for NCBI BLAST tools using the sequence of the distal ERG binding sites in mouse and human prostate cells. **(C)** Volcano plot for representative differentially expressed genes (DEGs) of RWPE1-ERG versus RWPE1-EV, up-regulated DEGs are marked by red, down-regulated DEGs are marked by blue. **(D)** Heatmap showing the expression of up-regulated luminal lineage genes and down-regulated basal lineage genes in RWPE1-ERG and RWPE1-EV cells respectively (RWPE1-ERG versus RWPE1-EV). **(E)** Box plots showing normalized counts (using DESeq2) of *ERG*, *KRT8*, *KRT18*, *TP63*, *KRT5* and *KRT14* respectively.
